# Oxidative stress induces inflammation of lens cells and triggers immune surveillance of ocular tissues

**DOI:** 10.1101/2021.10.16.464542

**Authors:** Brian Thompson, Emily A. Davidson, Ying Chen, David J. Orlicky, David C. Thompson, Vasilis Vasiliou

**Affiliations:** Department of Environmental Health Sciences, Yale School of Public Health, Yale University, 60 College Street, New Haven, CT, USA; Department of Cellular & Molecular Physiology, Yale School of Medicine, Yale University, New Haven, CT, USA; Department of Pathology, Anschutz School of Medicine, University of Colorado, Aurora, CO, USA; Department of Clinical Pharmacy, Skaggs School of Pharmacy and Pharmaceutical Sciences, University of Colorado Denver, Aurora, CO, USA

**Keywords:** Microphthalmia, Inflammation, Ocular Immune System, Glutathione, Lens, Oxidative Stress

## Abstract

Recent reports have challenged the notion that the lens is immune-privileged. However, these studies have not fully identified the molecular mechanism(s) that promote immune surveillance of the lens. Using a mouse model of targeted glutathione (GSH) deficiency in ocular surface tissues, we have investigated the role of oxidative stress in upregulating cytokine expression and promoting immune surveillance of the eye. RNA-sequencing of lenses from postnatal day (P) 1- aged *Gclc^f/f^;Le-Cre^Tg/−^* (KO) and *Gclc^f/f^;Le-Cre^−/−^* control (CON) mice revealed upregulation of many cytokines (e.g., CCL4, GDF15, CSF1) and immune response genes in the lenses of KO mice. The eyes of KO mice had a greater number of cells in the aqueous and vitreous humors at P1, P20 and P50 than age-matched CON and *Gclc^w/w^;Le-Cre^Tg/−^* (CRE) mice. Histological analyses revealed the presence of innate immune cells (i.e., macrophages, leukocytes) in ocular structures of the KO mice. At P20, the expression of cytokines and ROS content was higher in the lenses of KO mice than in those from age-matched CRE and CON mice, suggesting that oxidative stress may induce cytokine expression. *In vitro* administration of the oxidant, hydrogen peroxide, and the depletion of GSH (using buthionine sulfoximine (BSO)) in 21EM15 lens epithelial cells induced cytokine expression, an effect that was prevented by co-treatment of the cells with *N-*acetyl-L-cysteine (NAC), a antioxidant. The *in vivo* and *ex vivo* induction of cytokine expression by oxidative stress was associated with the expression of markers of epithelial-to-mesenchymal transition (EMT), α-SMA, in lens cells. Given that EMT of lens epithelial cells causes posterior capsule opacification (PCO), we propose that oxidative stress induces cytokine expression, EMT and the development of PCO in a positive feedback loop. Collectively these data indicate that oxidative stress induces inflammation of lens cells which promotes immune surveillance of ocular structures.

**Highlights:**

- Immune surveillance of ocular structures occurs in mouse eyes deficient in glutathione.
- Oxidative stress upregulates the expression of pro-inflammatory cytokines (e.g., GDF15, CSF1) in lens cells *in vitro* and *in vivo*.
- The upregulation of cytokines in lens cells is associated with markers of an epithelial-to-mesenchymal transition phenotype.
- Oxidative stress-induced inflammation and associated epithelial-to-mesenchymal transition may play a role in the development of posterior capsule opacification.

## Introduction

The lens, an avascular tissue surrounded by a thick basement membrane, has a unique relationship with the immune system. The lens is connected to the vascular and lymphatic systems throughout early eye development *via* the hyaloid vasculature, a temporary branch of the ophthalmic artery [1, 2]. Shortly after birth, the hyaloid vasculature regresses in a process mediated by macrophages [3, 4]. Notably, macrophages are also involved in the removal of apoptotic epithelial cells prior to lens cavity closure [5]. Following regression of the hyaloid vasculature and closure of the lens cavity, the lens has been thought to be immune-privileged, i.e., tolerant to the placement of allografts within the eye.

Several recent experimental results have challenged this dogma. First, the deletion of a cell-cell adhesion protein abundant in lens cells, N-cadherin, in the developing lens impairs lens development and induces immune surveillance of the lens and other ocular structures (i.e., cornea, vitreous humor, retina) from as early as embryonic day (E) 18.5 [6]. Second, following cataract surgery in mice, the expression of many cytokines is increased in lens epithelial cells prior to infiltration of immune cells into the remnant lens capsule [7]. Third, the zonule fibers that are connected to the lens provide a conduit for the trafficking of immune cells to the lens [6, 8, 9]. Fourth, damage to the cornea induces surveillance of the lens by immune cells [8]. Lastly, the chicken, mouse and human lens epithelium contain resident immune cells [9]. Despite this mounting evidence, the molecular mechanism(s) responsible for promoting immune surveillance of the lens remain to be elucidated.

Oxidative stress manifests as a result of an imbalance between antioxidants and oxidants (e.g., reactive oxygen species (ROS)) such that oxidants prevail [10]. Oxidative stress stimulates inflammation by activating the NF-κB signaling pathway [11, 12] and serving as a secondary messenger for the pro-inflammatory cytokine TNFα [13], thus ROS both stimulates and mediates inflammation. ROS is generated in the lens by a myriad of exogenous (e.g., radiation, pharmaceutical drugs, cigarette smoke) and endogenous (e.g., NADPH oxidases, cellular respiration) sources [14, 15], and may also be generated by infiltrating leukocytes [16]. Oxidative stress contributes to a common complication of cataract surgery, posterior capsule opacification (PCO) [17], which is characterized by the epithelial-to-mesenchymal transition of the lens epithelial cells that remain following cataract surgery [18–20].

In the present study, we describe that oxidative stress can induce an inflammatory response in lens epithelial cells. We report that the lens-specific deletion of *Glutamate-Cysteine Ligase Catalytic Subunit* (*Gclc*), a gene that encodes the rate-limiting enzyme in the biosynthesis of GSH, results in oxidative stress and triggers an inflammatory response that is characterized by a significant upregulation in the gene expression of several classes of cytokines in lens cells. We then replicated the system *in vitro* with cultured lens epithelial cells by treatment with buthionine sulfoximine (BSO), an irreversible glutamate cysteine ligase inhibitor, or hydrogen peroxide (H_2_O_2_) and found that this induces cytokine expression. Interestingly, treatment with BSO elicited the expression of more cytokines in these cells than did treatment with the oxidant H_2_O_2_, suggesting differential responses induced by these treatments. Lastly, we found that supplementation of lens epithelial explants with an antioxidant, *N-*acetyl-L-cysteine, reduced the expression of cytokines and prevented the induction of markers of EMT. Collectively, these results suggest that inflammation and PCO may both be prevented by post-cataract treatments that include an antioxidant or the upregulation of the endogenous antioxidant systems.

## Methods

### Mouse Lines

The creation of *Gclc* control (CON), *Gclc* knockout (KO) and *Le-Cre* control (CRE) mice used in this study have been previously described [21]. Briefly, *Gclc* homozygous floxed mice [22] and *Le-Cre* hemizygous mice [23] were crossed to delete *Gclc* from the cellular precursors of the eyelid, epithelium of the cornea, conjunctiva and lens from as early as embryonic day (E) 9. Mice of the three genotypes were maintained on a C57BL/6 and FVB/N mixed background. They were group-housed (no more than 5 mice per cage) and maintained on a 12-hour light-dark cycle, with food and water available *ad libitum.* All experiments were performed in strict accordance with the National Institutes of Health guidelines, and protocols were approved by the Yale University Institutional Animal Care and Use Committee.

### RNA sequencing (RNA-seq) Library Preparation and Sequencing

The RNA-seq data used in this study have been previously published [21]. Briefly, the lenses of mice aged postnatal day (P) 1 were collected from KO and CON mice (which were anesthetized by the isoflurane open drop method and euthanized by cervical dislocation) and stored in 200 μL RNAlater solution (Invitrogen, Waltham, MA) at −80°C until extraction. The lenses from three mice (i.e., six lenses) were pooled into a biological replicate and three biological replicates were used per genotype (i.e., 9 mice total per genotype). Total RNA was isolated from the biological replicates using the RNeasy Micro Kit (QIAGEN, Venlo, Netherlands) per the manufacturer’s instructions, and the RNA integrity number (RIN) was determined using the Agilent 2100 Bioanalyzer RNA 6000 Pico assay. cDNA libraries were prepared from total RNA samples with an RIN ≥ 8.0 using the NEBNext^®^ Single Cell/Low Input RNA Library Prep Kit for Illumina^®^ (New England BioLabs). An Illumina NovaSeq 6000 machine with an S4 flow cell was used to generate pairwise 100 bp reads (performed by the Yale Center for Genome Analysis).

### RNA-seq Analysis and Bioinformatics

The analysis of the RNA-seq data has been previously described in detail [21] and can be accessed at NCBI Gene Expression Omnibus (GEO) (accession number GSE175394). Briefly, all data analyses were performed using the Galaxy web platform [24] [accessed at usegalaxy.org], with default settings used for all tools (unless otherwise stated). Read sequence qualities were determined using the FastQC tool (v0.72+galaxy1), with low quality reads being trimmed using a sliding window (phred ≥ 20), and ambiguous bases (N) and any contaminating sequencing adapters removed using the Trimmomatic tool [25]. HISAT2 (v2.1.0+galaxy5)[26] was used to map the trimmed reads to the *Mus musculus* reference genome (GRCm38/mm10). featureCounts (v1.6.4+galaxy1) [27] was used to count mapped reads. DESeq2 (v2.11.0.6) [28] was used for differential expression analyses. Differentially-expressed genes (DEGs) were identified through satisfaction of the following criteria: ≥ ±1.0 log2 fold change (log2FC) and adjusted P value < 0.05 (Benjamini-Hochberg method[29]).

The Database for Annotation, Visualization and Integrated Discovery (DAVID) bioinformatics resource [30] was used to perform gene ontology (GO) functional annotation analysis on identified DEGs. Ingenuity Pathway Analysis (IPA) (Version 52912811, Ingenuity Systems, QIAGEN) was used to identify ‘canonical pathways’ in the upregulated DEGs. The cytokine-cytokine receptor interaction pathway map was generated using the KEGG Mapper program (31423653).

### Histological Analysis of Mouse Eyes

P1-aged mice were euthanized by swift decapitation and a piece of the tail was removed for genotyping by PCR, as previously described [22]. Mice aged P20 and P50 were anesthetized by the isoflurane open drop method, euthanized by cervical dislocation. The eyes were then rapidly enucleated and any contaminating tissues were removed. The mouse heads or eyes were fixed in Davidson’s Solution for 24 hours at 4°C and subsequently stored in 70% EtOH. Yale Pathology Tissue Services (YPTS) processed the tissues for histological analysis, i.e., conducted paraffin embedding, sectioning (5 μm thickness) and mounting onto glass slides. YPTS then either stained the resultant slides with hematoxylin and eosin (H&E) or subjected them to immunohistochemical analysis (per their standard protocols). At least two H&E-stained sections from the eyes of three CON, CRE or KO mice (aged P1, P20 or P50) were imaged with a Nikon Eclipse E200 microscope with an Axiocam 503 camera (Zeiss) attached and the number of cells within the aqueous and vitreous humors were counted using the NIH Image J software [31]. Cell counts are presented as means and associated standard deviation.

### Cell Culture

The mouse lens epithelial cell line, 21EM15, was obtained from Dr. Salil Lachke (Department of Biological Sciences, University of Delaware). Cells were cultured in Dulbecco’s Modified Eagle Medium (DMEM) (Thermo Fisher Scientific Inc., MA) supplemented with 10% fetal bovine serum (Sigma-Aldrich, MO), 1% antibiotic-antimycotic (Sigma-Aldrich, St. Louis, MO) and 1% MEM non-essential amino acids (Sigma-Aldrich, MO) in 60 mm dishes (Corning, NY) in a humidified atmosphere of 5% CO_2_ in air at 37°C. Cells were treated with 500 μM BSO (Sigma-Aldrich, MO) for 48 hours or with 25 μM hydrogen peroxide (Cole Palmer, IL) for 24 hours to induce oxidative stress [32, 33]. In other experiments, cells were concomitantly treated with 10 mM *N-*acetyl-L-cysteine (Sigma-Aldrich, MO) and 500 μM BSO for 48 hours. At the end of the treatment period, cells were rinsed once with phosphate buffered saline (PBS, Gibco, MA) and removed from the culture dish by a 3-minute treatment with 0.05% trypsin (Gibco, MA) and the trypsin neutralized with equal parts cell culture medium. The disassociated cells were transferred to a 1.5 mL microcentrifuge tube (Eppendorf, Hamburg, Germany) and a cell pellet was generated by centrifugation at 500 g for 3 min at room temperature. The cell culture media was aspirated from the cell pellet and the pellet was washed twice with room temperature PBS by gently resuspending the cell pellet in PBS, centrifugation at 500 g for 3 min at room temperature, and aspiration of the PBS. After the final wash, the cell pellet was stored in 100 μL of PBS at −80°C for latter analysis.

### Lens Epithelial Explant Establishment

*Gclc^w/w^* mice aged P20 were used for the establishment of lens epithelial explants, as previously described [34, 35]. Briefly, mice were anesthetized, euthanized and their eyes enucleated as described in the histological analysis section (above). The lenses were removed and placed into a 35 mm culture dish (Corning, NY) containing pre-warmed (37°C) Medium 199 supplemented with 0.1% fetal bovine serum (Sigma-Aldrich, MO), 1% antibiotic-antimycotic (Sigma-Aldrich, St. Louis, MO) or Medium 199 supplemented with 0.1% fetal bovine serum (Sigma-Aldrich, MO), 1% antibiotic-antimycotic (Sigma-Aldrich, St. Louis, MO), and 10 mM NAC (Thermo Fisher Scientific). A hole was made at the posterior pole, the lens capsule opened, and the fiber cells gently removed. The lens capsule was pinned to the bottom of the culture dish such that the adherent epithelial cells were exposed to the medium. Explants were then individually cultured in a humidified atmosphere of 5% CO_2_ at 37°C for 24 hours. Six explants were pooled to make one sample and 3 pooled samples were used for each experimental condition (hence a total of 18 mice were used per condition).

### RNA Isolation and RT-qPCR

#### RNA isolation from mouse lenses

Three CON, CRE and KO mice aged P20 were anesthetized by the isoflurane open drop method, euthanized by cervical dislocation and lenses dissected. Dissected lenses were immediately placed in 100 μL RNAlater solution (Thermo Fisher Scientific), flash frozen in liquid nitrogen and stored at −80° C until processing. Total RNA was isolated from the mouse lenses using the RNeasy Plus Micro Kit (QIAGEN, Venlo, Netherlands) per the manufacturer’s instructions. Briefly, the lenses were removed from the RNAlater solution, placed in 350 μL RLT Plus buffer (QIAGEN, Venlo, Netherlands), and total RNA was isolated per the manufacturer’s instructions.

#### RNA isolation from 21EM15 cells

Total RNA was isolated from the 21EM15 cells using the RNeasy Plus Mini Kit (QIAGEN, Venlo, Netherlands) per the manufacturer’s instructions. Each harvested cell pellet was resuspended in 600 μL RLT Plus Buffer and were lysed with a Tissuelyser (QIAGEN, Venlo, Netherlands) at a frequency of 30 Hz for 2 min at 4°C. Total RNA was isolated per the manufacturers instructions.

#### RNA isolation from lens epithelial explants

Lens epithelial explants (6 explants per experiment per condition) from three independent experiments (i.e., 18 explants total per experimental condition) were placed in were placed in 100 μL RNAlater solution, flash frozen in liquid nitrogen and stored at −80°C until processing. Total RNA was isolated from these explants using the RNeasy Plus Micro Kit (QIAGEN, Venlo, Netherlands) per the manufacturer’s instructions.

### RT-qPCR

Each total RNA sample was quantified and analyzed for purity using a spectrophotometer (Nanodrop ND-1000). Five hundred ng of total RNA (from 21EM15 cells or mouse lenses) or 10 ng of total RNA (from explants) were reverse transcribed using the iScript cDNa Synthesis Kit (Bio-Rad, CA) per the manufacturer’s instructions. One ng of cDNA from the explants was then pre-amplified using the Quantabio PerfeCTa PreAmp Supermix (Quantabio, MA) with the same primers as used for qPCR (per the manufacturer’s instructions) and diluted 20-fold. Ten ng of cDNA from the 21EM15 cells, 1 uL of preamplified cDNA from the explants or 75 ng of cDNA from the lenses was then used to estimate the abundance of specific mRNA transcripts using the iTaq Universal SYBR Green Supermix (Bio-Rad, CA) on a CFX96 Real-Time PCR System (Bio-Rad, CA). Relative mRNA transcript abundance was estimated using the ΔCt method [36] with the housekeeping gene, GAPDH, used as an internal normalization control for each sample. Primer sequences used are provided in Supplemental Table 1.

### Analysis of ROS levels

ROS levels were assayed as previously described [37]. Briefly, three P20-aged CON, CRE and KO mice were anesthetized by the isoflurane open drop method, euthanized by cervical dislocation and lenses dissected. The two freshly isolated lenses from each animal were placed into a single 96-well plate containing 200 μL Medium 199 (Sigma-Aldrich) maintained at 4°C. Dihydrorhodamine 123 (DHR) (7.5 μM) (Invitrogen, MA), a colorless stain that easily passes through membranes and is oxidized by ROS into rhodamine 123, and 1 drop of NucBlue Live Ready Probe (Hoescht 33342, Thermo Fisher Scientific, MA) were added to each well and incubated at 4°C for 30 minutes. Stained lenses were washed three times in 4°C Medium 199 and then 200 μL 4°C PBS (Gibco, MA) was added to each well prior to measuring DHR fluorescence intensity at Ex/Em of 507/529 nm and Hoescht 33342 at Ex/Em of 360/460 nm with a microplate reader (Spectramax M3, Molecular Devices). DHR relative fluorescence units (RFU) was expressed as a ratio of the Hoescht 33342 RFU in the same tissue. 1EM15 cells were cultured in a 60 mm culture dishes (Corning, NY) and treated with either 500 μM BSO, 25 μM H_2_O_2_ or 500 μM BSO + 10 mM NAC (for the periods described above). Lens epithelial explants were cultured in normal culture media or media containing 10 mM NAC for 24 or 48 hours, respectively. At the end of the treatment period, the culture medium was aspirated and replaced by 3 mL of ice-cold Medium 199. Dihydrorhodamine 123 (7.5 μM) and 5 drops of NucBlue Live Ready Probe (Hoescht 33342, Thermo Fischer Scientific, MA) were added to each dish and allowed to incubate for 30 minutes at 4°C. Stained cells/explants were then washed three times in 4°C Medium 199 and finally 3mL of fresh 4°C Medium 199 was added to each dish. The cells or explants were then either imaged on a AxioVert.A1 microscope (Zeiss) with an Axiocam 305 camera (Zeiss) and using a Photoflor LM 75 light source (89 North, VT).

### Western Blot Analysis

21EM15 cells were cultured in 60 mm cell culture dishes (Corning, NY), treated with 500 μM BSO or an equivalent volume of medium (control) for 48 hours, harvested by treatment with 0.05% trypsin for 3 min and the trypsin was neutralized with equal parts cell culture medium. The dissociated cells were transferred to a 1.5 mL microcentrifuge tube (Eppendorf, Hamburg, Germany) and subjected to centrifugation at 500 g for 3 min at room temperature. The cells were lysed by resuspension of the pellet in 250 μL RIPA buffer (1% Nonidet P40, 0.5% sodium deoxycholate, 0.1% SDS in PBS), incubation for 10 mins on ice and passing the suspension 10 times each through 22, 25 and 28 gauge series of needles (in that order) [38]. The protein samples were then subjected to centrifugation at 14,000 rpm for 10 min at 4°C and the supernatant was collected. Protein concentrations in the supernatant were quantified using the Pierce BCA Protein Assay Kit (Thermo Fisher Scientific, MA) according to the manufacturer’s instructions. Thirty μg of supernatant protein was resolved on a 4-20% SDS-PAGE gradient gel (Bio-Rad, CA) and transferred to a 0.2 μm nitrocellulose blot (Bio-Rad, CA). Primary antibodies (1:1000) directed against NF-κB P65 (Cell Signaling Technologies, 8242T), P-NF-κB P65 (Cell Signaling Technologies, 3033T), IKK-β (Cell Signaling Technologies, 8943S), IκBα (Cell Signaling Technologies, 4814T), P-IκBα (Cell Signaling Technologies, 2859T) or GAPDH (Abcam, ab9485) were used for immunoblotting. Horse radish peroxidase-conjugated goat anti-rabbit or goat anti-mouse secondary antibodies (1:5000, Cell Signaling Technologies, 7074P2) were used to visualize immunolabeled proteins. Quantitation of band densities was performed using NIH Image J software [31]. Target protein expression was normalized to the corresponding GAPDH expression or unphosphorylated protein, as appropriate. Data are presented as the mean density (and associated standard deviation) of the normalized protein.

### Statistical Analysis

Differences between gene expression and protein expression were determined using Student’s unpaired t-test or one-way ANOVA with *post-hoc* Dunnett’s test correction. Differences between cell counts were determined using one-way ANOVA with *post-hoc* Dunnett’s test correction. Differences in ROS are expressed as fold change (of CON) and significance was determined using a one-way ANOVA with *post-hoc* Dunnett’s test correction. All statistical analyses were conducted using GraphPad Prism version 9.1.1 for PC, GraphPad Software, La Jolla California, USA. P < 0.05 was considered significant.

## Results

### *Gclc* deletion induces an inflammatory response in the lenses of neonatal KO mice

We have previously described the *Gclc^f/f^;Le-Cre^Tg/−^* knockout (KO) mouse model used in this study [21]. Briefly, *Gclc* gene was specifically deleted from surface ectoderm-derived tissues (i.e., corneal epithelium, conjunctiva, eyelid, lens) from as early as embryonic day (E) 9 by crossing *Gclc^f/f^* mice [22] with *Le-Cre* transgene mice [23]. KO mice have an overt microphthalmia phenotype that is characterized by vacuolation of the lens fiber cells at birth, hypercellularity of the retina, cornea and iris by P20, and severe retinal infolding by P50 (Supp. Fig. 1). Controlling for the *Le-Cre* transgene [39], revealed that the microphthalmia phenotype and morphological changes in the KO mice are distinct from those in *Le-Cre* transgene hemizygous mice, termed CRE (Supp. Fig. 3).

We have previously described the impaired lens development phenotype of KO mice by conducting RNA-seq analysis on lens tissue from KO and CON mice aged P1 [21]. Fifty-three genes associated with the Gene Ontology (GO) term “immune system process” were upregulated in the lenses of KO mice relative to those in CON mice (Fig. 1A, red dots). Complete analysis of the GO terms overrepresented among the up-regulated genes in the lenses of KO mice revealed many terms associated with the immune system/inflammation (e.g., “immune system process”, “inflammatory response”, “neutrophil chemotaxis”, “chemotaxis”, “cell adhesion”, “chemokine-mediated signaling pathway”, “positive regulation of inflammatory response”, “immune response”) (Fig. 1B). As a sensitivity analysis, the upregulated genes in KO mice were also analyzed with Ingenuity Pathway Analysis; several canonical pathways associated with the immune system/inflammation were overrepresented among the upregulated genes in KO mice (i.e., “agranulocyte adhesion and diapedesis”, “granulocyte adhesion and diapedesis”, “hepatic fibrosis/hepatic stellate cell activation”, “dendritic cell maturation”, “atherosclerosis signaling”, “phagosome formation”, “neuroinflammation signaling pathway”, “TREM1 signaling”, “IL-10 signaling”) (Fig. 1C). Mapping the upregulated transcripts onto the Kyoto Encyclopedia of Genes and Genomes (KEGG) Cytokine-Cytokine Receptor Interaction pathway revealed 42 of the genes involved in this pathway were upregulated (Fig. 1D, highlighted in red) in the lenses of KO mice belonged to the CC subfamily, CXC subfamily, γ-chain utilizing, IL4-like, IL6/12-like, IL10/28-like, Interferon family, IL1-like cytokines, TNF family, and TGF-β family. Of the 45 differentially expressed genes (DEGs) in the lenses of KO mice (aged P1) that mapped to the Cytokine-Cytokine Receptor Interaction pathway, only three genes (*Ccl27*, *Cnftr, 4-1Bbl)* were downregulated (Fig. 1D, highlighted in blue). Furthermore, many cytokines were among the top 25 upregulated genes in the lenses of KO mice (i.e., *Ccl7, Ccl4, Gdf15, Cxcl16*) (Supp. Table. 2).

**Figure 1.**
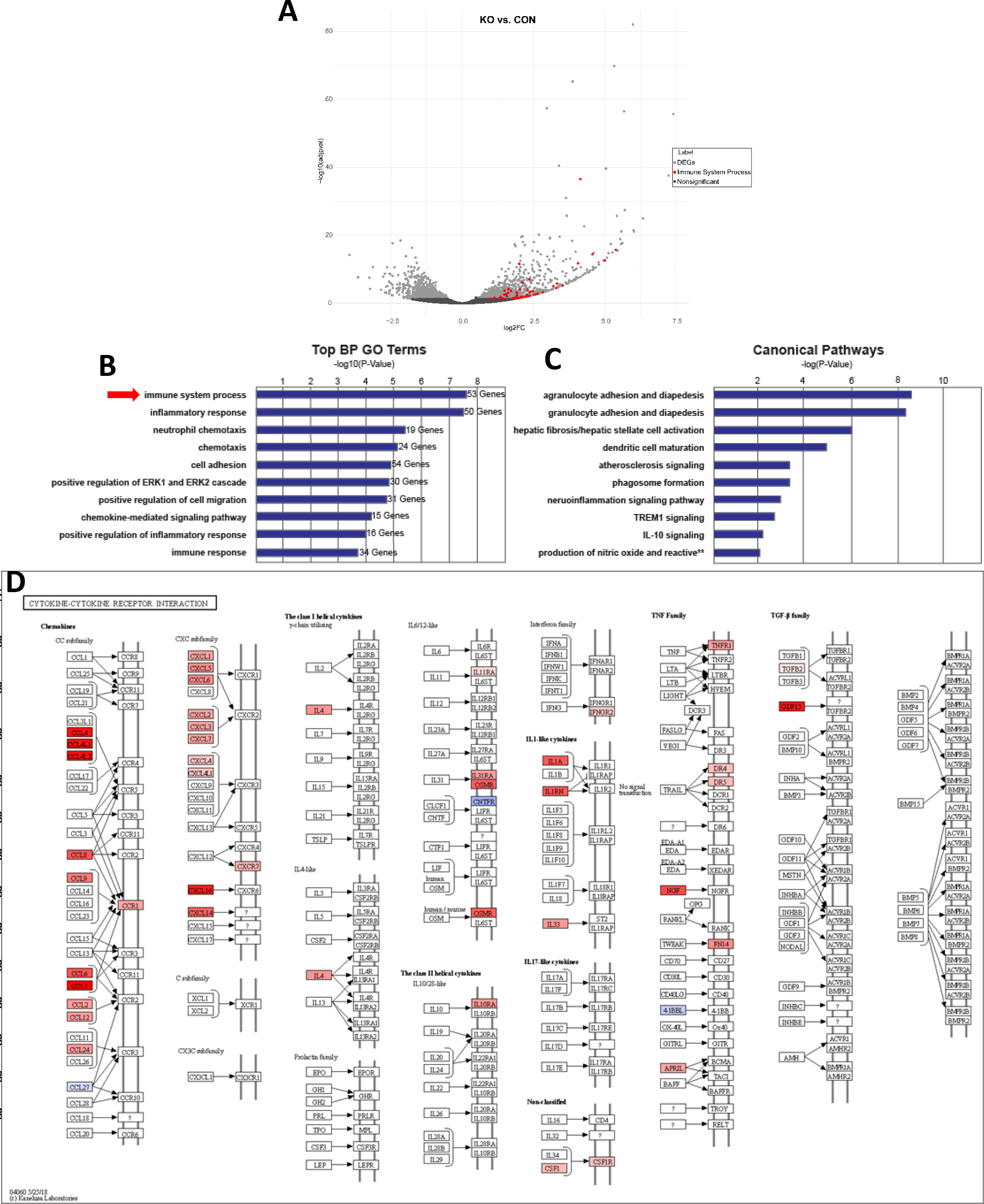
Inflammatory response in the lenses of KO mice at postnatal day 1. RNA-sequencing of postnatal day (P) 1 lenses revealed changes in genes in KO mice relative to CO mice. (**A**) Volcano plot illustrating the 530 downregulated (P<0.05) and 1022 upregulated (P<0.05) genes in KO mice (pooled samples from 3 mice). The 53 upregulated DEGs that represent the biological processes GO term “immune system process” (red arrow) are indicated by red dots. The probability (−log_10_(adjusted P)) (Y-axis) is Benjamini-Hochberg-corrected. Fold changes (KO vs. CON) are displayed on the X-axis as log_2_FC. **B**) Top biological processes (BP) gene ontology (GO) terms among upregulated genes. The −log10(P-Value) for each term is indicated by a blue bar. The number of DEGs among each term is indicated at the righthand end of each bar. P-values are Benjamini-Hochberg adjusted. (**C**) Top canonical pathways among upregulated genes as identified with Ingenuity Pathway Analysis. The −log10(P-Value) for each canonical pathway is indicated by a blue bar. P-values are Benjamini-Hochberg adjusted. (**D**) The Kyoto Encyclopedia of Genes and Genomes (KEGG) Mapper Cytokine-Cytokine Receptor Interaction map annotated to highlight DEGs. Colors: blue shading, downregulated; red shading, upregulated. Darker shading colors indicate greater differential expression.

### Innate immune cells infiltrate ocular structures of KO mice

Morphological analysis of the eyes from KO mice showed an increased presence of cells in the aqueous and vitreous humors at P1 compared with age-matched CON mice (Supp. Figs. 1B, 1H; Supp. Fig. 2). Immunohistochemical staining revealed that the vitreous humor of KO and CON mice at P1 had cells of leukocytic origin present, as indicated by both CD45- (Fig. 2A-C, J-L, open arrows) and CD11b- (Fig, 2D-F, M-O, arrowheads) staining cells; some CD45-positive cells appeared to cross the lens capsule in the eyes of KO mice (Fig. 2L, open arrows). Immunohistochemical staining for CD68 indicated the presence of macrophages in the vitreous humor of both KO and CON mice at P1 (Fig. 2G-I, P-S, closed arrows), which was expected given the role of macrophages in regression of the hyaloid vasculature [3, 4]. As anticipated, the lenses from CRE mice aged P1 also had a similar number of cells in the vitreous humor as age-matched CON mice (Supp. Fig. 2). Immunohistochemical staining revealed that some of the cells found in the vitreous humor of CRE mice by H&E staining (Supp. Fig. 3A (B’) were CD45- (Supp. Fig. 3B (A’, B’, arrowheads) or CD68-staining (Supp. Fig. 3B (F’, closed arrows)).

**Figure 2:**
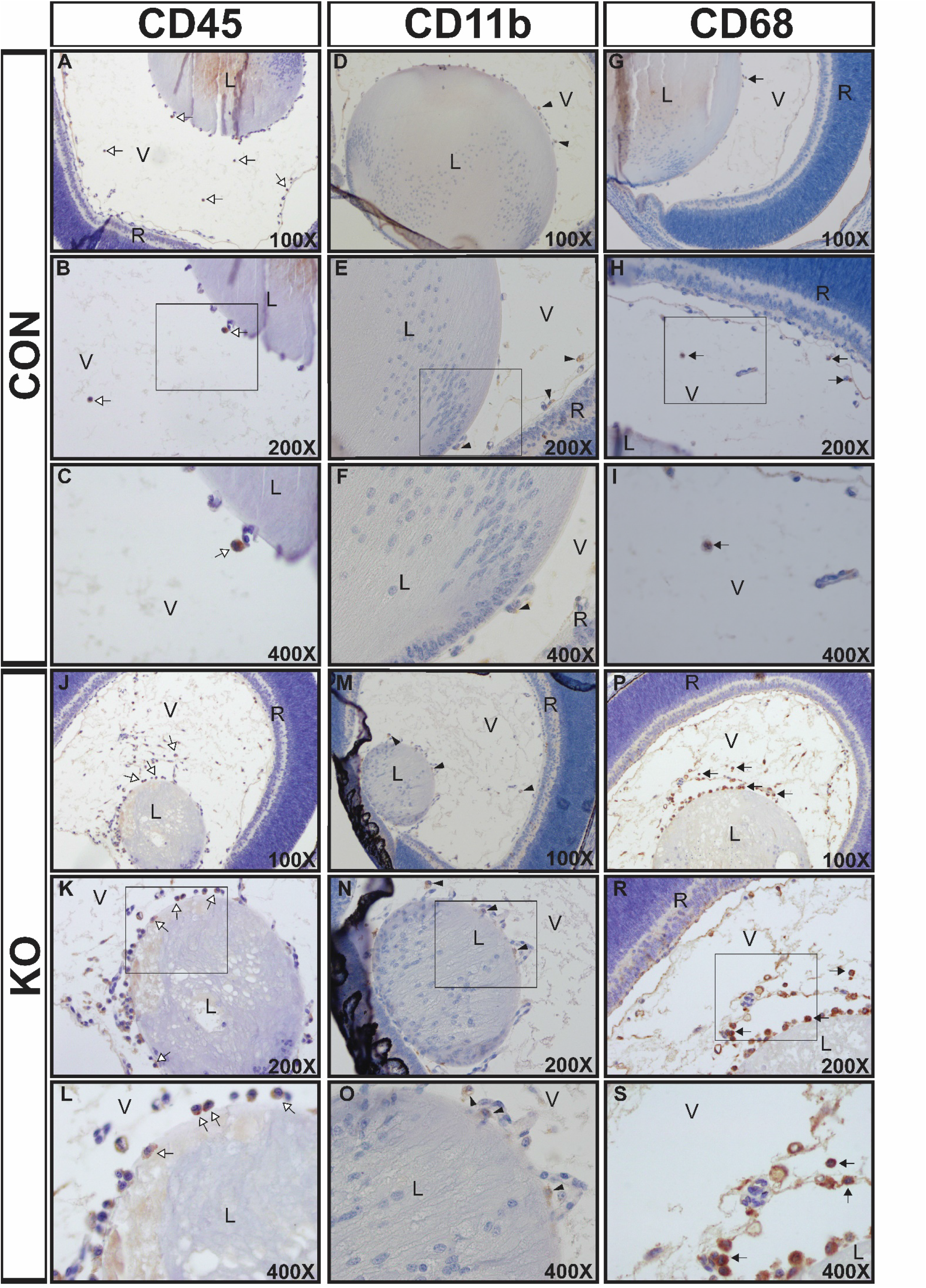
Immunohistochemical analysis of immune cells in the eyes of KO and CON mice aged postnatal day 1. Eyes from CON (**A-I**) and KO (**J-S**) mice aged postnatal day (P) 1 were subjected to immunohistochemical staining and counterstained with hematoxylin. CD45 (**A-C**, **J-L**, open arrows) and CD11b (**D-F**, **M-O**, arrowheads) staining marks non-macrophage leukocytes. CD68 staining (**G-I**, **P-S**, arrows) marks macrophages. Regions in squares in panels D, E, H, K, N and R are shown at higher magnification in **C**, **F**, **I**, **L**, **O** and **S**, respectively. Abbreviations: C, cornea; L, lens; R, retina; V, vitreous humor. Magnification is indicated in the lower right corner of each image.

Histological analysis of the eyes from CON mice aged P20 revealed that the aqueous and vitreous humors were almost entirely devoid of cells (Supp. Figs. 1C, 1D; Supp. Fig. 2). In contrast, histological analysis of the eyes from KO mice aged P20 revealed an increase in the number of cells in the aqueous and vitreous humor (Supp. Figs. 1I, 1J; Supp. Fig. 2). Eyes from KO mice aged P20 had CD45- (Fig. 3A-C, open arrows) and CD11b- (Fig. 3D-F, arrowheads) staining cells of leukocytic origin and CD68 staining macrophages present in the vitreous humor (Fig. 3G-I, closed arrows). As previously noted [6], lens malformation can induce immune surveillance of many ocular structures, a phenomena that was observed in the eyes of KO mice (Fig. 3). In the eyes of KO mice at P20, CD11b-staining cells were present in the retina, aqueous humor and corneal epithelium (Fig. 3E, F, arrow heads); CD45-staining cells appeared to be exiting the retina into the vitreous humor (Fig. 3C, open arrows); CD68-staining cells were found in the aqueous humor in the KO mice that had formed an aqueous humor (Fig. 3H, closed arrows). Histological analysis of CRE mice aged P20 revealed the presence of an elevated number of cells in the vitreous humor compared with CON mice (Supp. Fig. 2; Supp. Fig. 3A (C’, D’)); KO mice aged P20 had an elevated number of cells in the vitreous humor compared with CRE mice (Supp. Fig. 2; Supp. Fig. 3A (C’, D’)). Immunohistochemical analysis of these eyes revealed CD45-staining cells in the corneal endothelium (Supp. Fig. 3B (D’, closed arrow)) and CD68 staining cells in the vitreous humor (Supp. Fig. 3B (G’, H’, open arrow)). As expected, the eyes from CON and CRE mice aged P50 were almost completely devoid of cells in the aqueous and vitreous humors (Supp. Fig. 1E, F, closed arrows; Supp. Fig. 2; Supp. Fig. 3A (E’, F’)), whereas the eyes from KO mice aged P50 had a greater number of cells in the aqueous and vitreous humors compared with the eyes of CON mice (Supp. Fig. 1K, M; Supp. Fig. 2).

**Figure 3:**
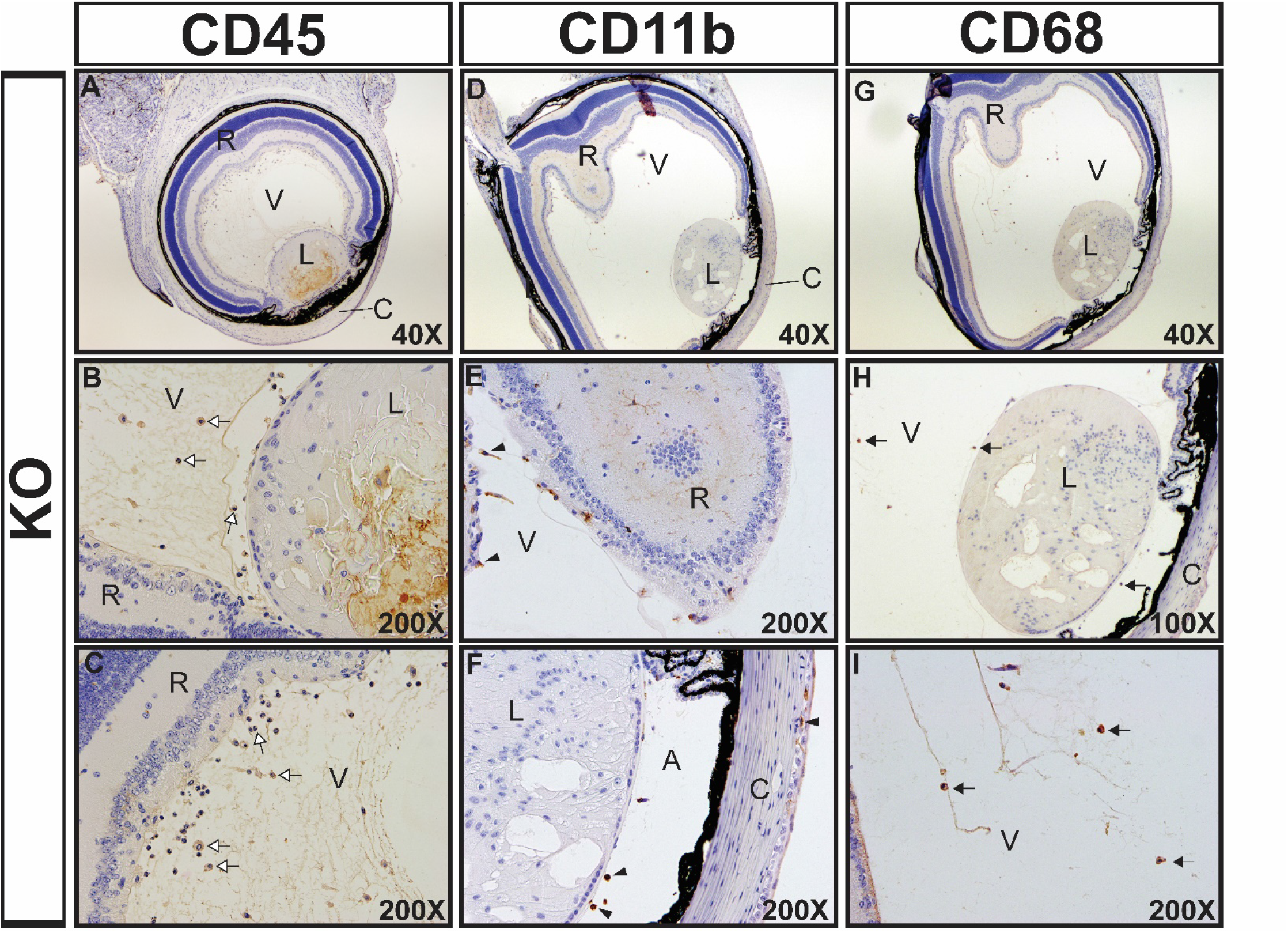
Immunohistochemical analysis of immune cells in the eyes of KO mice aged postnatal day 21. Eyes from KO mice aged postnatal day (P) 20 were subjected to immunohistochemical staining and counterstained with hematoxylin. CD45- (**A-C**, open arrows) and CD11b staining (**D-F**, arrowheads) marks non-macrophage leukocytes. CD68-staining (**G-I**, arrows) marks macrophages. Abbreviations: A, aqueous humor C, cornea; L, lens; R, retina; V, vitreous humor. Magnification is indicated in the lower right corner of each image.

### *Gclc* deletion induces expression of immune system related genes and causes oxidative stress in the lenses of KO mice

Given the alterations in immune surveillance in the eyes of KO mice, the expression of cytokine genes in the lenses of CON, KO and CRE mice aged P20 were evaluated (Fig. 4A). Since the expression of some cytokine genes in the lenses of CON mice were undetectable by RT-qPCR, the gene expression is displayed as ΔCt, such that a lower ΔCt corresponds to a greater gene expression. For genes with detectable expression levels in the lenses of CON mice (*Ccl7*, *Ccl2*, *Gdf15*), only the lenses of KO mice had an increase in gene expression (compared with CON mice) (Fig. 4A). The expression of *Ptprc* and *Cxcl16* were both upregulated in the lenses of KO mice compared with CRE mice (Fig. 4A). Levels of ROS were elevated in the lenses of P20-aged KO mice relative to similarly aged CON and CRE mice (Fig. 4B).

**Figure 4:**
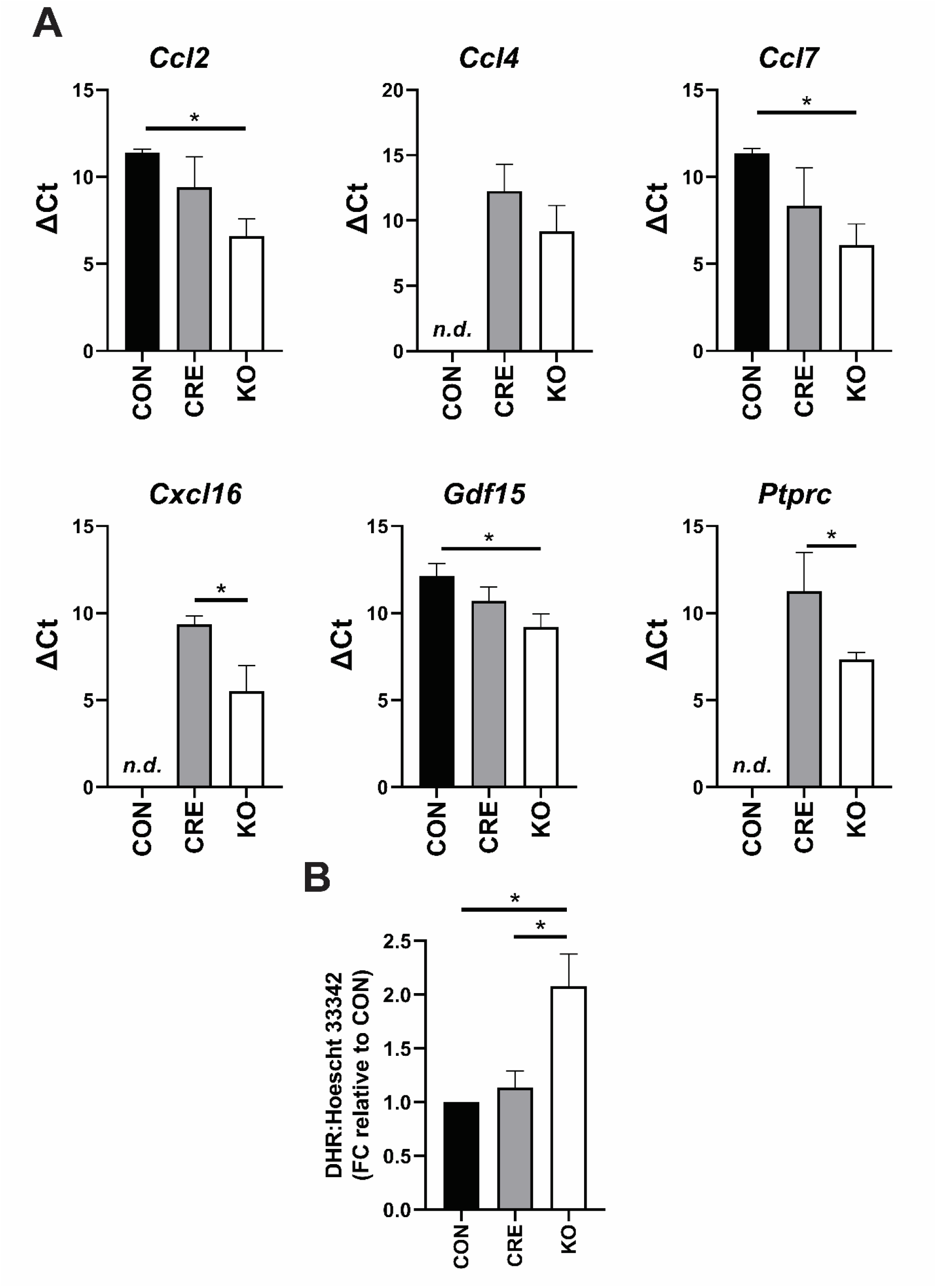
Cytokine gene expression and ROS levels in the lenses of CON, KO and CRE mice aged postnatal day 20. (**A**) Lenses from CON, CRE and KO mice aged postnatal (P) 20 were analyzed for induction of genes involved in the inflammatory response by reverse transcriptase (RT) quantitative PCR (qPCR), as calculated by the ΔCt method. GAPDH was used as an internal normalization control. Gene expression (ΔCt) is presented as the mean and associated standard deviation from 3 mice. * P < 0.05, one-way ANOVA with *post-hoc* Dunnett’s test, compared to group indicated by horizontal bar. *n.d.* = not detectable (**B**) ROS production in the lenses of CON, KO and CRE mice aged P20. ROS levels were monitored using the ROS probe dihydrorhodamine 123 (DHR) and cell nuclei were labelled with Hoescht 33342. DHR relative fluorescence units (RFU) was expressed as a ratio of the Hoescht 33342 RFU in the same tissue. Data are presented as fold change (FC) relative to CON with associated standard deviation. * P < 0.05, one-way ANOVA t-test with *post-hoc* Dunnett’s test, compared to genotype indicated.

### Oxidative stress upregulates cytokine expression in lens epithelial cells

To further evaluate the influence of oxidative stress on cytokine expression in the lens, 21EM15 mouse lens epithelial cells were depleted of GSH by treatment with buthionine sulfoximine (BSO), a chemical inhibitor of GSH biosynthesis. This intervention induced an oxidative stress response as indicated by the upregulation of the antioxidant response element (ARE) genes [40], *Gclc* and *Hmox1* (Fig. 5A), and a marked increase in ROS formation (Fig. 5D). Since 21EM15 cells do not express all of the cytokines that were differentially expressed in the lenses of KO mice [41] (Fig. 1), the effect of BSO-induced oxidative stress on the expression of cytokines could only be evaluated for a limited number of cytokines. BSO-induced oxidative stress upregulated the expression of *Cxcl1*, *Cxcl12*, *Gdf15*, *Ccl7*, *Ccl2*, and *Csf1* (Fig. 5A). Similarly, treatment of 21EM15 cells with 25 μM hydrogen peroxide (H_2_O_2_) for 24 hours induced an oxidative stress response that involved upregulated expression of *Gclc* and *Hmox1* (Fig. 5B) and increased ROS formation (Fig. 5D). The H_2_O_2_ treatment upregulated the expression of the cytokines *Ccl2*, *Ccl7*, and *Cxcl1* but failed to induce expression of the cytokines *Cxcl12, Csf1* or *Gdf15* (Fig. 5B). Co-treatment of 21EM15 cells for 48 hours with BSO and the antioxidant *N-*acetyl-L-cysteine (NAC) prevented the induction of oxidative stress by BSO as indicated by no induction of the expression of *Gclc* and *Hmox1* and no increase in ROS formation (Figs. 5C & D). The NAC co-treatment also prevented induction of the cytokines *Ccl7, Cxcl12, Gdf15, Ccl2, Csf1,* and *Cxcl1* (Fig. 5C). Collectively, these results suggest that oxidative stress is capable of inducing cytokine expression in lens epithelial cells.

**Figure 5.**
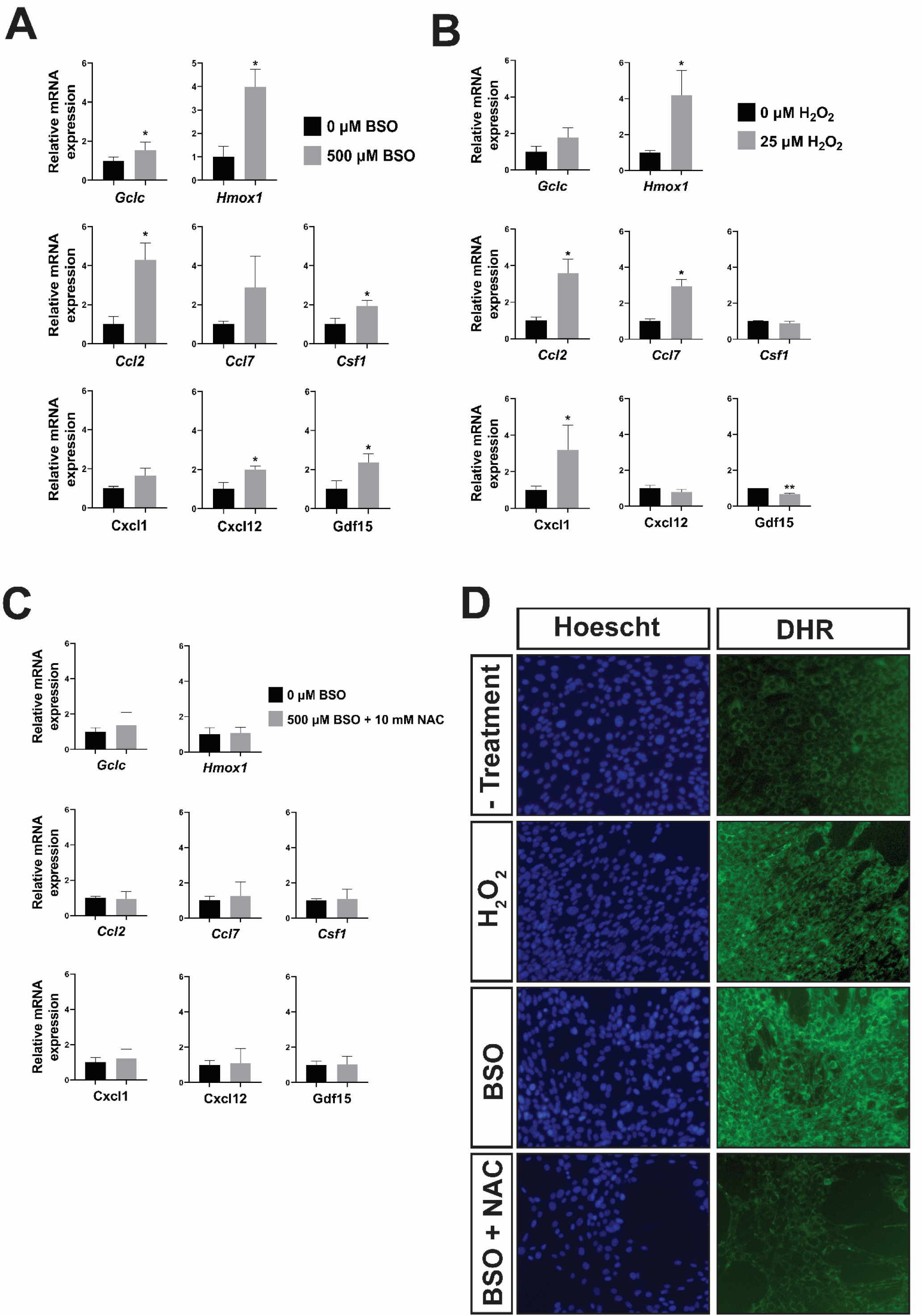
Oxidative stress induces cytokine expression in lens epithelial cells. Lens epithelial cells (21EM15) were treated with (**A**) 500 μM buthionine sulfoximine (BSO) for 48 hours, (**B**) 25 μM H_2_O_2_ for 24 hours, or (**C**) 500 μM BSO and 10 mM *N-*acetyl-L-cysteine (NAC) for 48 hours. Gene expression was determined by reverse transcriptase (RT) quantitative PCR (qPCR), as calculated by the ΔCt method. GAPDH was used as an internal normalization control. Gene expression is displayed as the average of the fold change relative to control and associated standard deviation from three independent experiments. * P < 0.05 Student’s unpaired t-test, compared to untreated (0 μM) cells. (**D**) ROS production was determined in 21EM15 cells treated with 500 μM BSO for 48 hours, 25 μM H_2_O_2_ for 24 hours, or 500 μM BSO + 10 mM NAC for 48 hours using the ROS probe dihydrorhodamine 123 (DHR). Cell nuclei were labelled with Hoescht 33342. Images were taken with the same camera settings and magnification (200x).

Given that oxidative stress in lens epithelial cells can activate NF-κB [11, 42–44], an inducer of cytokine gene expression [45], the activation of NF-κB in cultured lens epithelial cells and *in vivo* lens was evaluated (Supp. Fig. 4). NF-κB activation can be mediated through several mechanisms: i) phosphorphorlyation of NF-κB (P65 subunit), ii) proteasomal degradation of NF-κB, iii) proteasomal degradation of IKK-β and/or iv) phosphorylation of IκBα [45]. The effect of BSO-induced oxidative stress (in 21EM15 cells) on these mechanisms for NF-κB activation were evaluated by Western blot (Supp. Figs. 4A & B). BSO treatment failed to influence NF-κB activation (Supp. Figs. 4A & B). Expression of genes induced by activated NF-κB in lens cells [46] (i.e., *Birc5, Bcl2, Bcl2l1, Birc2*) were also unchanged in cells treated with BSO (Supp. Fig. 4C). NF-κB-activated genes were similarly not induced in the lenses of KO mice aged P1 (Supp. Fig 4D).

### Epithelial-mesenchymal transition is associated with the increased cytokine expression

Oxidative stress and, in particular, low GSH-induced oxidative stress transforms lens epithelial cells to myofibroblasts [17]. This transformation is a feature of epithelial-mesenchymal transition (EMT) [18–20], a process characterized by increased expression of alpha-smooth muscle actin (α-SMA) [19]. Therefore, we investigated if the increases in cytokine gene expression were associated with an EMT phenotype (Fig. 6). α-SMA positive cells were found throughout the lens epithelium from the anterior pole to the posterior pole in the lenses of KO mice aged P1 (Fig. 6A (B’, C’, closed arrow)). Analysis of α-SMA expression in the lenses of KO mice aged P20 similarly revealed α-SMA positive cells throughout the lens epithelium (Fig. 6A (E’,F’, closed arrow)). Intriguingly, the lens epithelium of CRE mice also possessed α-SMA staining cells, although this phenotype was not present in the lenses of all CRE mice (in contrast to KO mice where it was present in all eyes analyzed)) (Supp. Figs. 5 A, B & F, arrow).

**Figure 6:**
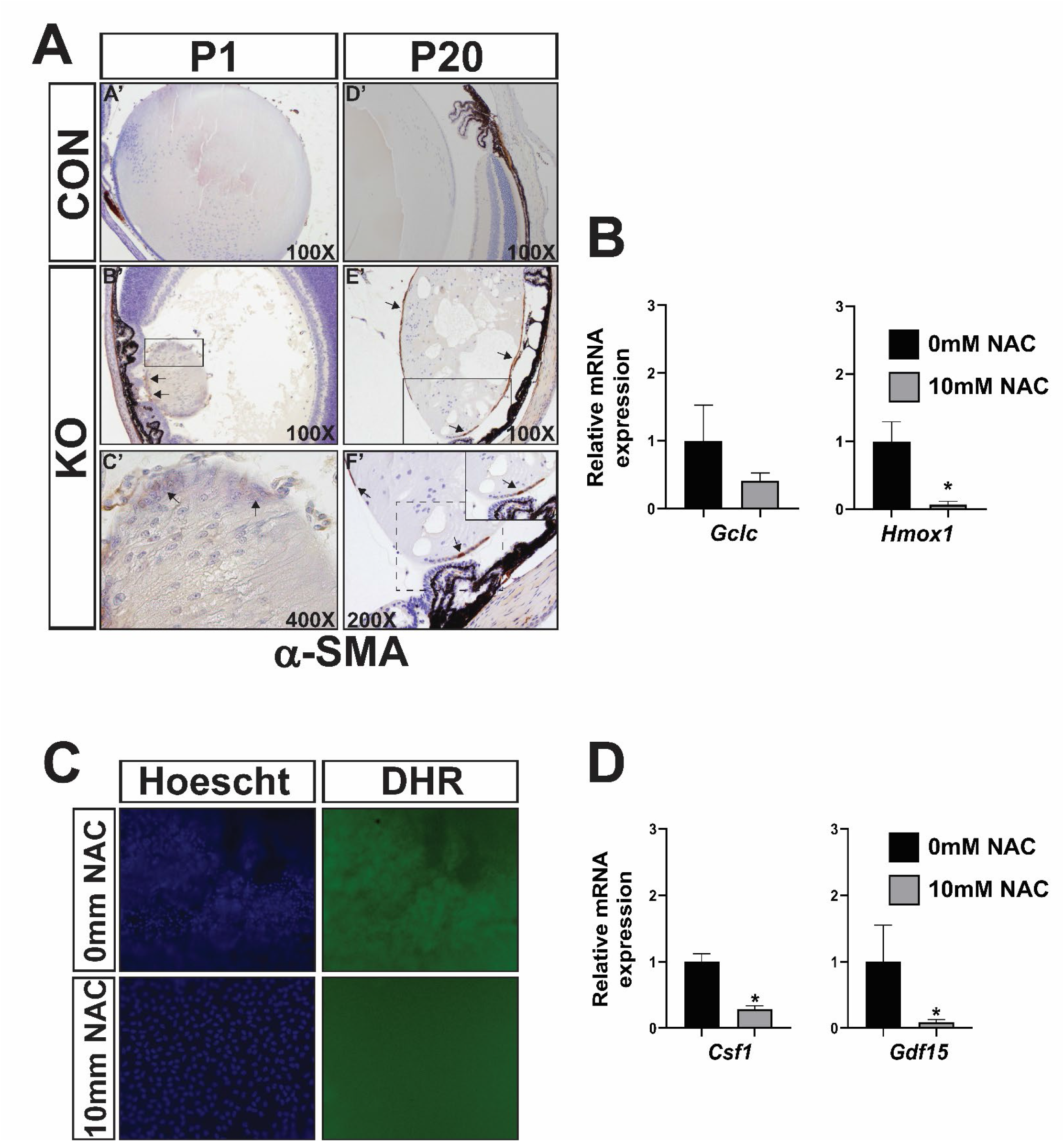
Analysis of markers of EMT in KO lenses aged P20 and detection of markers of oxidative stress, and cytokine expression in lens epithelial explants. (**A**) Immunohistochemical staining for alpha-smooth muscle actin (α-SMA) and counterstaining with hematoxylin in the eyes of KO mice aged postnatal day (P) 1 and 20. Positive staining for α-SMA (closed arrows) indicates cells undergoing EMT. Regions demarcated by boxes in B’ and E’ are shown in higher magnification in C’ and F’, respectively. Dashed box in F’ shown at 400X magnification in inset. Magnification is indicated in the lower right corner of each image. (**B**-**E**) Lens epithelial explant systems were established in normal media (0mM NAC) or media containing 10 mM *N-*acetyl-L-cysteine (10mM NAC) and cultured for 24 hours. (**B**) Expression of antioxidant response element genes. (**C**) ROS levels were monitored by dihydrorhodamine 123 (DHR) staining. Cell nuclei were labelled with Hoescht 33342. Images were taken with the same camera settings and magnification (200x). NAC- = 0mM NAC. (**D**) Expression of cytokines. Gene expression was determined by reverse transcriptase (RT) quantitative PCR (qPCR), as calculated by the ΔCt method. GAPDH was used as an internal normalization control. Gene expression is presented as the mean of the fold change relative to control (0 μM NAC) and standard deviation. * P < 0.05, Student’s unpaired t-test, compared to 0mM NAC.

### Mitigation of oxidative stress in lens epithelial explants prevents induction of cytokine expression

Given the links between oxidative stress, cytokine expression and EMT [9, 17, 47–50], we wished to explore if antioxidant administration could prevent the induction of cytokines and an EMT phenotype in a lens cataract surgery model, viz. the lens epithelial explant model. This model is a powerful tool for understanding lens epithelial cell biology and pathology, including posterior capsule opacification (PCO), a common complication of cataract surgery [35]. A recent RNA-seq experiment found that lens epithelial explants upregulate the expression of cytokines, antioxidant response element genes and markers of EMT within 24 hours of being established (personal communication with Dr. Michael Robinson, [51]). Thus, we investigated the role of oxidative stress in these processes. Treatment of lens epithelial explants with 10 mM NAC reduced oxidative stress as evidenced by reduced expression of *Gclc* and *Hmox1* (Fig. 6B) and reduced the presence of ROS (Fig. 6C). The treatment of lens epithelial explants with 10 mM NAC only had a non-significant effect on preventing EMT, as evidenced by the trend towards increased expression of the lens identity genes, *Cdh1* and *Maf*, and decreased expression of known markers of EMT, *Acta2* (Supp. Fig. 6). The NAC treatment also prevented the induction of *Csf1* and *Gdf15* (Fig. 6E), but did not prevent the induction of *Ccl2*, *Ccl7*, or *Cxcl1* (Supp. Fig. 7).

## Discussion

Mounting evidence challenges the long-held notion that the lens is isolated from the immune system [6–9], which may have important implications for ocular health. The mechanism(s) by which immune surveillance of the lens and other ocular structures can be triggered by damage to the lens remains to be understood. The upregulation of cytokine expression in human lens epithelial (HLE-B3, SRA01/04) cells by H_2_O_2_ [32] and ultraviolet B radiation [52] suggests that oxidative stress may elicit an inflammatory response in the lens. To our knowledge, a detailed characterization of oxidative stress-induced lens inflammation has yet to be performed *in vitro* or *in vivo*. In the present study, we have extended our previous unexpected finding that deletion of *Gclc* from the developing lens (and resulting oxidative stress) upregulates the expression of 42 cytokines and promotes immune surveillance of the cornea, aqueous humor and vitreous humor [21]. In addition, we also report that oxidative stress can induce cytokine expression in cultured lens epithelial cells (*Ccl2, Ccl7, Csf1, Cxcl1, Cxcl12, Gdf15)* and lens epithelial explants (*Csf1, Gdf15)*. Lastly, our data suggest that the upregulation of cytokines is associated with EMT.

We found that oxidative stress upregulated the expression of 42 cytokines in the neonatal lenses of *Gclc* KO mice and that immune cells were present in the cornea, retina, aqueous humor and vitreous humor of these mice. Cytokines are small secreted proteins that modulate cell growth and differentiation, and activation and migration of immune cells to areas of tissue damage. Chemokines are a class of cytokine secreted from cells that recruit immune cells to a site of tissue damage [53]. Interestingly, amongst the upregulated genes in the lenses of KO mice were potent chemokines for many leukocytes, e.g., *Ccl6* [54], *Ccl4* [55], *Ccl7* [56], *Ccl2* [57], *Cxcl12* [58]. It is reasonable to expect that increased secretion of these cytokines contributed to the observed recruitment of the immune cells to the eyes of KO mice (as detected by immunohistochemical staining for CD45, CD11b and CD68). The induction of these chemokines in lens epithelial cells (21EM15) by oxidative stress strongly suggests that the increased expression of these genes *in vivo* is largely due to increased expression in lens epithelial cells rather than from infiltrating immune cells. This contention is supported by the finding that many of the genes encoding these chemokines are poised for expression (in a state of “open” chromatin) [59], expressed in lens epithelial cells *in vitro* [41] and *in vivo* [60, 61], and rapidly upregulated in mouse lens epithelial cells following mock cataract surgery [7]. Collectively, these data are consistent with oxidative stress upregulating cytokine expression in lens epithelial cells, leading to enhanced immune surveillance of ocular structures. Thus, mitigation of oxidative stress and/or cytokine expression in lens epithelial cells may dampen immune surveillance of ocular structures.

There are many mechanisms by which oxidative stress can induce cytokine expression in cells, one of which is activation of NF-κB [11, 42–44]. The results derived from lens epithelial (21EM15) cells and lenses from KO mice do not support a mechanistic role of NF-κB activation in our experimental settings. Other mechanisms by which oxidative stress can induce cytokine expression include altered histone code [62–66], damaged mitochondrial DNA [67, 68], upregulated IRF1 expression [69] and activated AP-1 [66, 70], STAT3 [71], NLRP3 [72, 73] and MAPK [32] signaling pathways. Which of these mechanisms mediates the effects in the KO lens remains to be established.

Inflammation manifesting in ocular structures after cataract surgery is thought to result from disruption of the blood-aqueous barrier that is formed by the iris and ciliary, prostaglandin release from the iris and ciliary body, or lens-induced uveitis [74–76]. Our results raise the possibility that oxidative stress-induced inflammation of lens epithelial cells may contribute to the inflammation after cataract surgery. This proposal is supported by the finding that, in humans, markers of oxidative stress (i.e., malondialdehyde) and cytokines are elevated in the aqueous humor after cataract surgery [74]. Given that: i) some researchers consider a common complication of cataract surgery, posterior capsule opacification (PCO), to be a form of postoperative inflammation [77], ii) myofibroblasts produce cytokines [78], iii) chemokine gene expression is higher in fibroblasts than in epithelium [79] and iv) lens epithelial cells undergoing EMT express chemokines [80], we hypothesize that the increased expression of cytokine genes induced by oxidative stress may be related to EMT of lens epithelial cells. This hypothesis is supported by our *ex vivo* experiments in which the treatment of lens epithelial cell explants with the antioxidant NAC reduced oxidative stress, attenuated the development of EMT markers, and prevented the induction of the chemokines *Csf1* and *Gdf15*. It has been reported that ultraviolet B radiation can induce the expression of *Gdf15* in lens epithelial cells [52] and *Gdf15* expression promotes EMT in colorectal cancers [47]. Our findings that *Gdf15* was among the 25 most upregulated genes in the lenses of KO mice and was associated with the development of markers of EMT led us to speculate that oxidative stress-induced expression of *Gdf15* may be involved in EMT of lens epithelial cells during PCO. However, our current data cannot eliminate the possibility that the induction of *Csf1* may also be involved in the pathogenesis of PCO and/or fibrosis of the lens, as suggested by a report that found a role for CSF1 in macrophage-induced fibrosis of the lens [81]. Future studies will be needed to discern these possibilities.

A hallmark of EMT in the lens has been the presence of α-SMA positive cells, which arise from the lens epithelial cells [82]. However, recent work has demonstrated that α-SMA expressing cells in the lens can arise from multiple sources including: i) populations of G8 positive mesenchymal precursor cells that reside within the lens epithelium [83], ii) resident lens immune cells [9, 50], and iii) infiltrating immune cells [6] (including macrophages [81]). Given the extensive damage to the lens in KO mice and recruitment of immune cells to the lens, it is likely that the source of α-SMA positive cells is not homogenous and may involve some (or all) of the described cellular sources.

Taking into account our data and numerous previous reports [9, 17, 47–50], we postulate that a positive feedback loop exists wherein increases in oxidative stress and cytokine gene expression are connected in a manner that ultimately results in PCO (Fig. 7). Such a positive feedback loop may have important implications for postoperative cataract surgery care.

**Figure 7:**
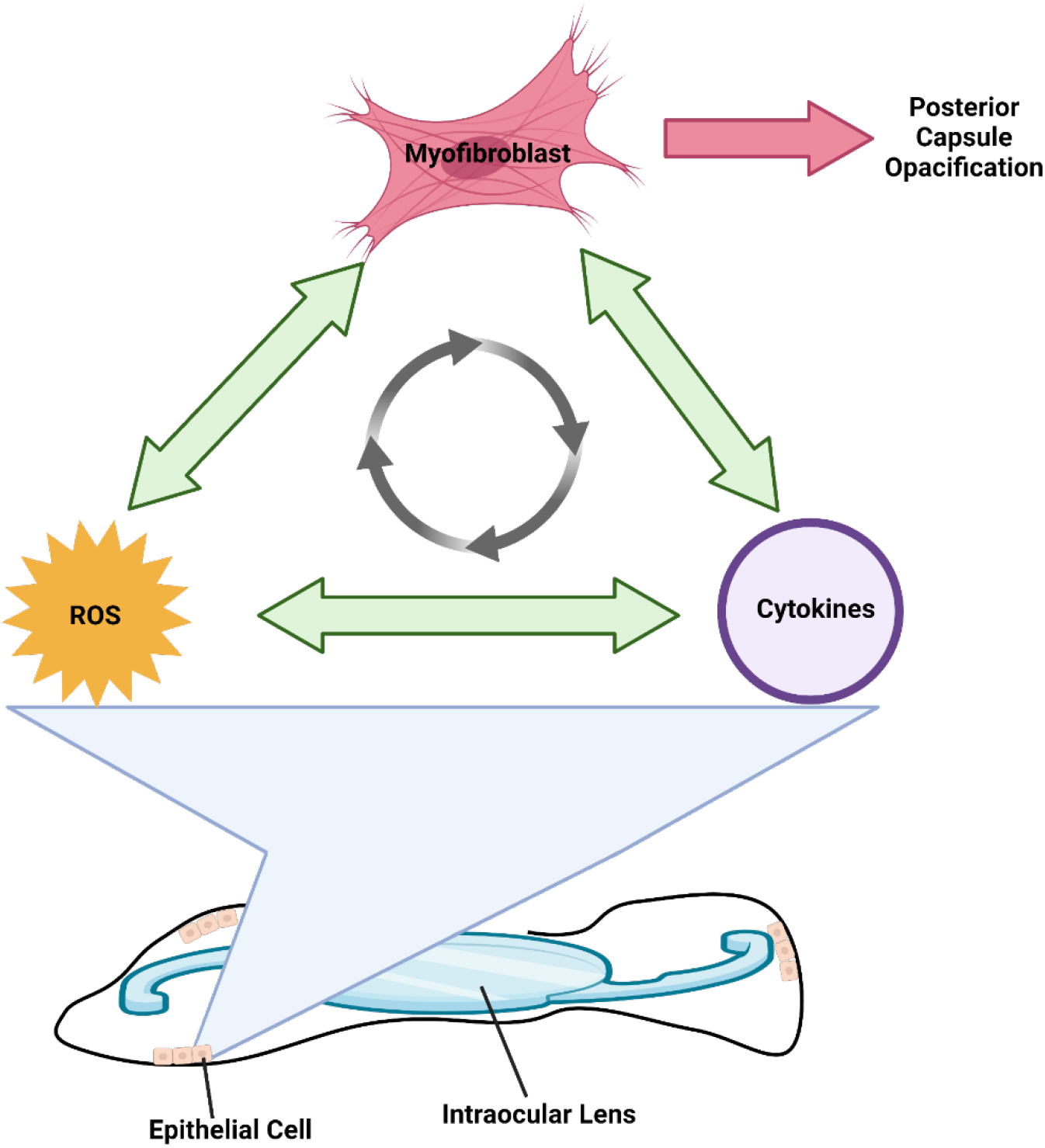
Proposed positive feedback loop by which oxidative stress and cytokine expression promote Posterior Capsule Opacification (PCO). We propose that a positive feedback loop reinforces the development of PCO. Increased ROS-driven oxidative stress and/or increased cytokine expression not only, induce the transdifferentiation of the remaining epithelial cells into myofibroblasts (EMT) but, increase each other and together reinforce EMT. Consequently, as more of the remaining lens epithelial cells undergo EMT occurs, PCO develops. Created with bioender.com.

Inflammation following cataract surgery is typically controlled by the combined administration of corticosteroids and non-steroidal anti-inflammatory drugs [76]. While this treatment strategy is effective at dampening the inflammation, its ability to prevent PCO is equivocal [84–89]. The failure of this treatment strategy to prevent PCO may be due to its inability to mitigate oxidative stress [90–93], which is essential for preventing the development of PCO-relevant physiological and molecular events [17, 32, 49, 94]. It is hoped that the results of the present study will motivate the exploration of treatment strategies that include antioxidant agents as a means to further reduce postoperative inflammation and prevent PCO formation. Should such a strategy be effective, it may reduce the immense burden of PCO which can affect upwards of 25% of adults [95] who undergo cataract surgery.

## Acknowledgements

We would like the thank Mr. Rolando Garcia-Milan for his advice on the RNA-seq analysis, the laboratory of Dr. Mark Petrash for their assistance with the histology and the members of the laboratory of Dr. Michael Robinson for their thoughtful discussions.

## Author Contributions

V.V. conceived of the study. B.T., Y.C., and V.V. designed the experiments. B.T. performed the experiments. B.T., E.A.D., Y.C., D.J.O., D.C.T. and V.V. analyzed data, discussed results, wrote and edited the manuscript.

## Supplemental Figures

**Supplemental Figure 1:**
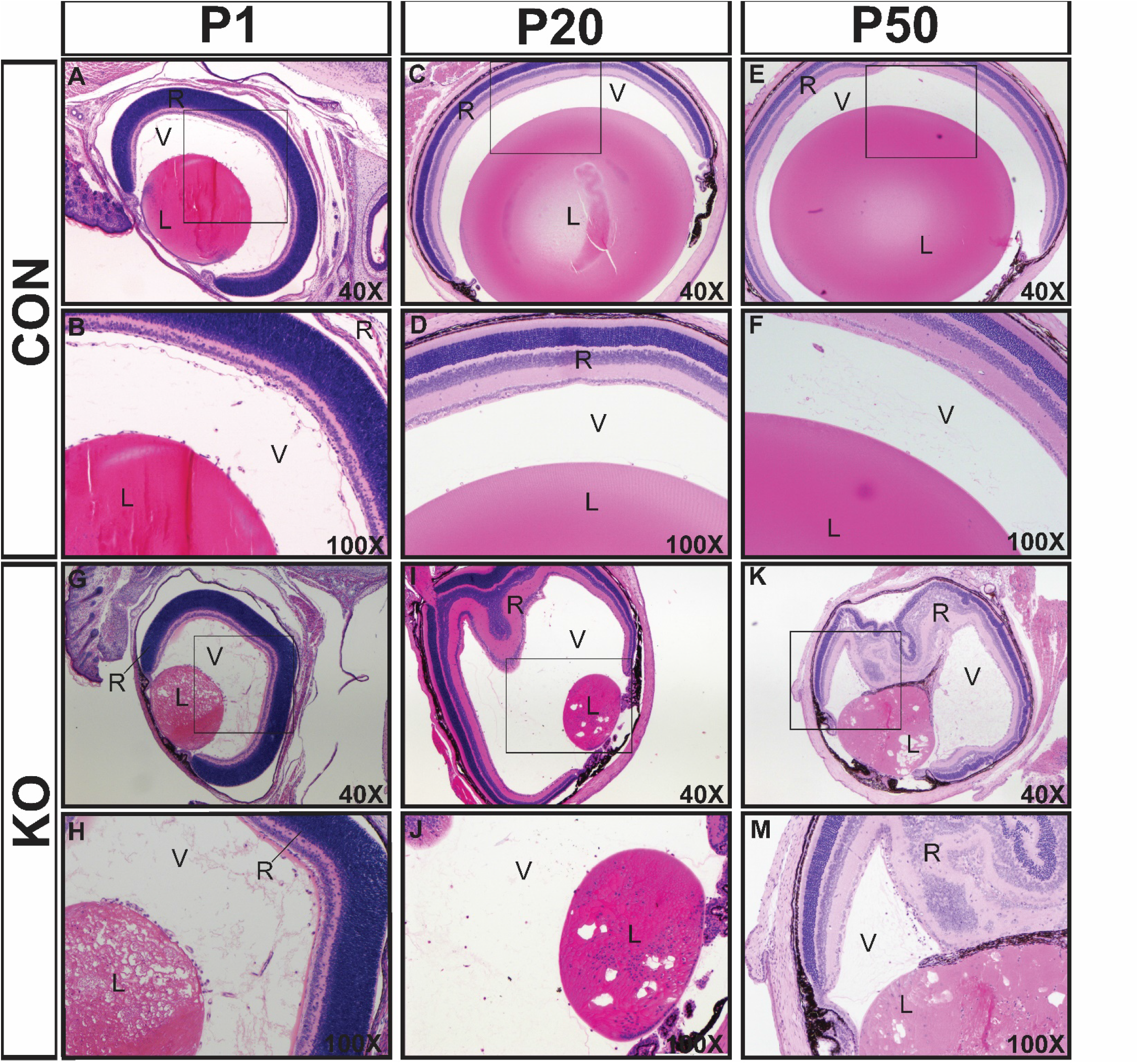
Histological analysis of eyes from CON and KO mice. Tissue sections of eyes from CON and KO mice aged postnatal day (P) 1, 20 and 50 were analyzed for changes in gross morphology. Hematoxylin & eosin staining of eyes from CON (A-F) and KO (G-M) mice. Abbreviations: C, cornea; L, lens; R, retina; V, vitreous humor. Regions demarcated by boxes in A, C and E are shown in higher magnification in B, D and F, respectively. Magnification is indicated in the lower right corner of each image.

**Supplemental Figure 2:**
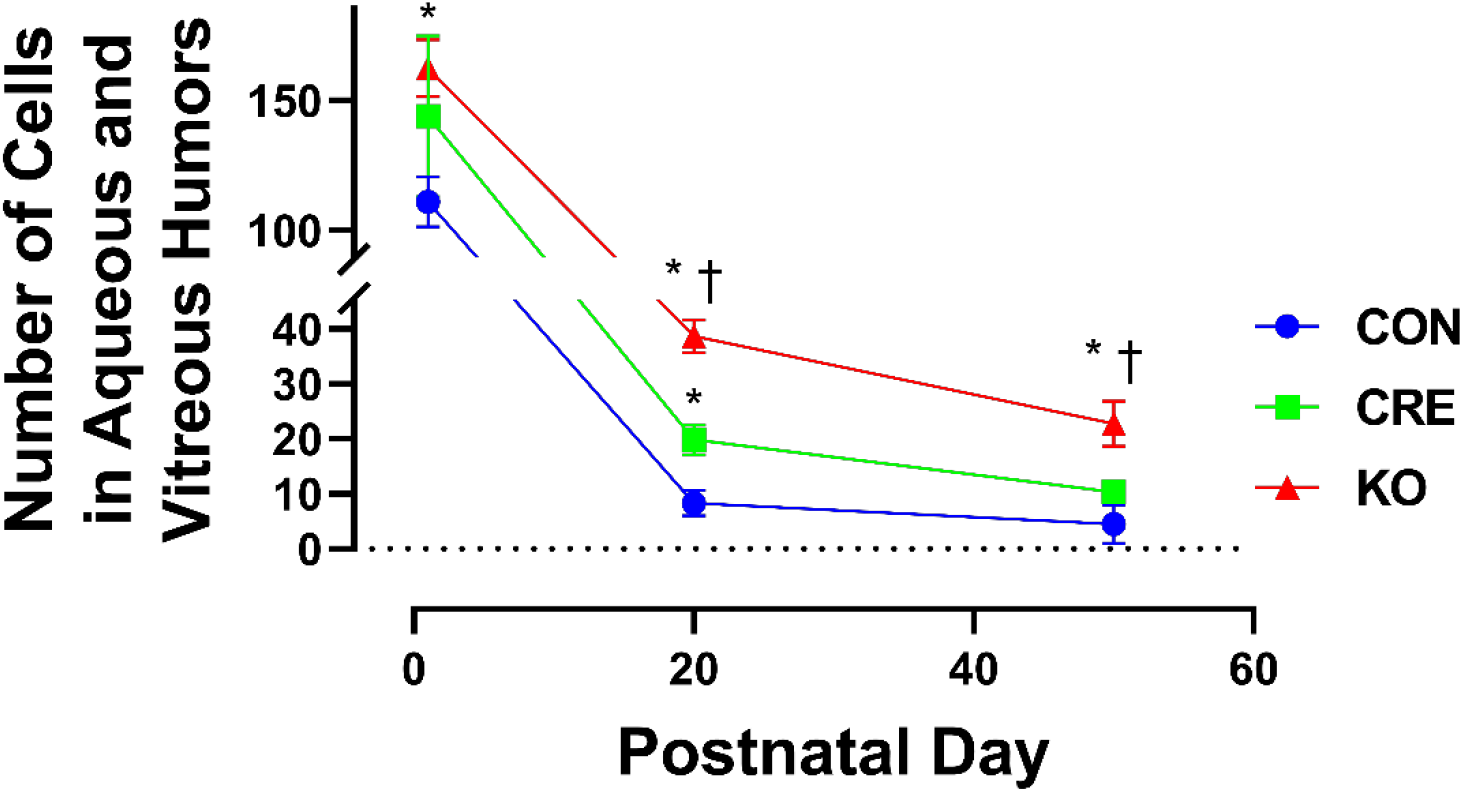
Cell counts in the Aqueous and Vitreous Humors. The eyes from CON, CRE and KO mice aged P1, P20 or P50 (n=3) were sectioned, stained by H&E and the number of cells in aqueous and vitreous humor counted. Data are presented as the mean ± SD from 3 mice (the number of cells in the pair of eyes from each mouse were averaged). * P < 0.05, ANOVA with *post-hoc* Tukey’s test correction, compared with CON; †, P < 0.05, ANOVA with *post-hoc* Tukey’s test correction, compared with CRE

**Supplemental Figure 3:**
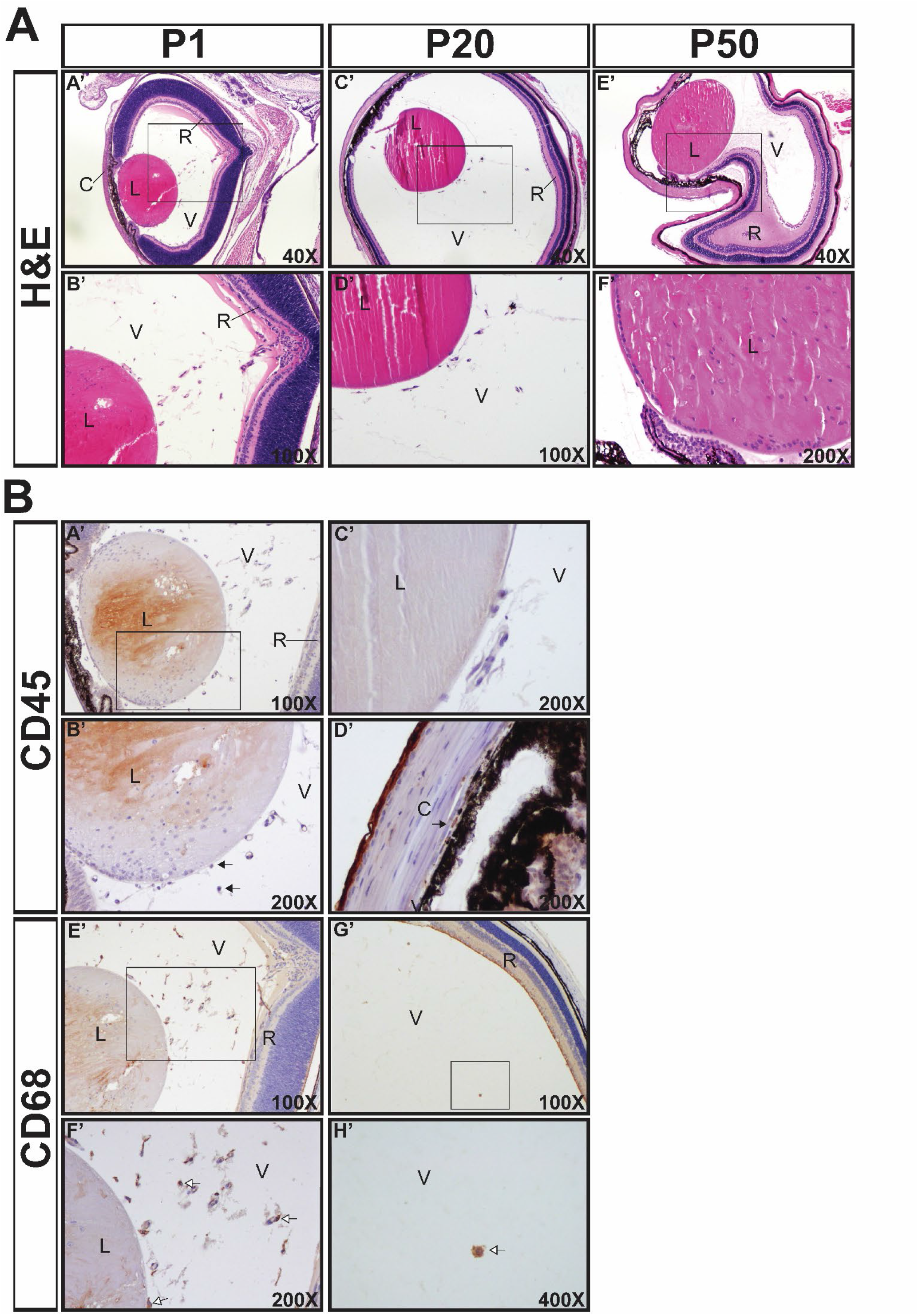
Characterization of immune surveillance of ocular structures in the eyes of CRE mice. The eyes from CRE mice aged postnatal day (P) 1, 21 and 50 were analyzed for the presence of cells in ocular structures. (**A**) Hematoxylin & eosin (H&E) staining at P1 (A’, B’), P21 (C’, D’) and P50. (E’ F’). (**B**) Immunohistochemical staining against CD45 in P1 (A’, B’) and P21 (C’, D’) CRE mice. Closed arrows indicate leukocytes. Immunohistochemical staining against CD68 in P1 (E’, F’) and P21 (G’, H’) CRE mice. Open arrows indicate macrophages. Regions demarcated by boxes in A’, E’ and G’ are shown in higher magnification in B’, F’ and H’, respectively. Abbreviations: C, cornea; L, lens; R, retina; V, vitreous humor. Magnification is indicated in the lower right corner of each image.

**Supplemental Figure 4:**
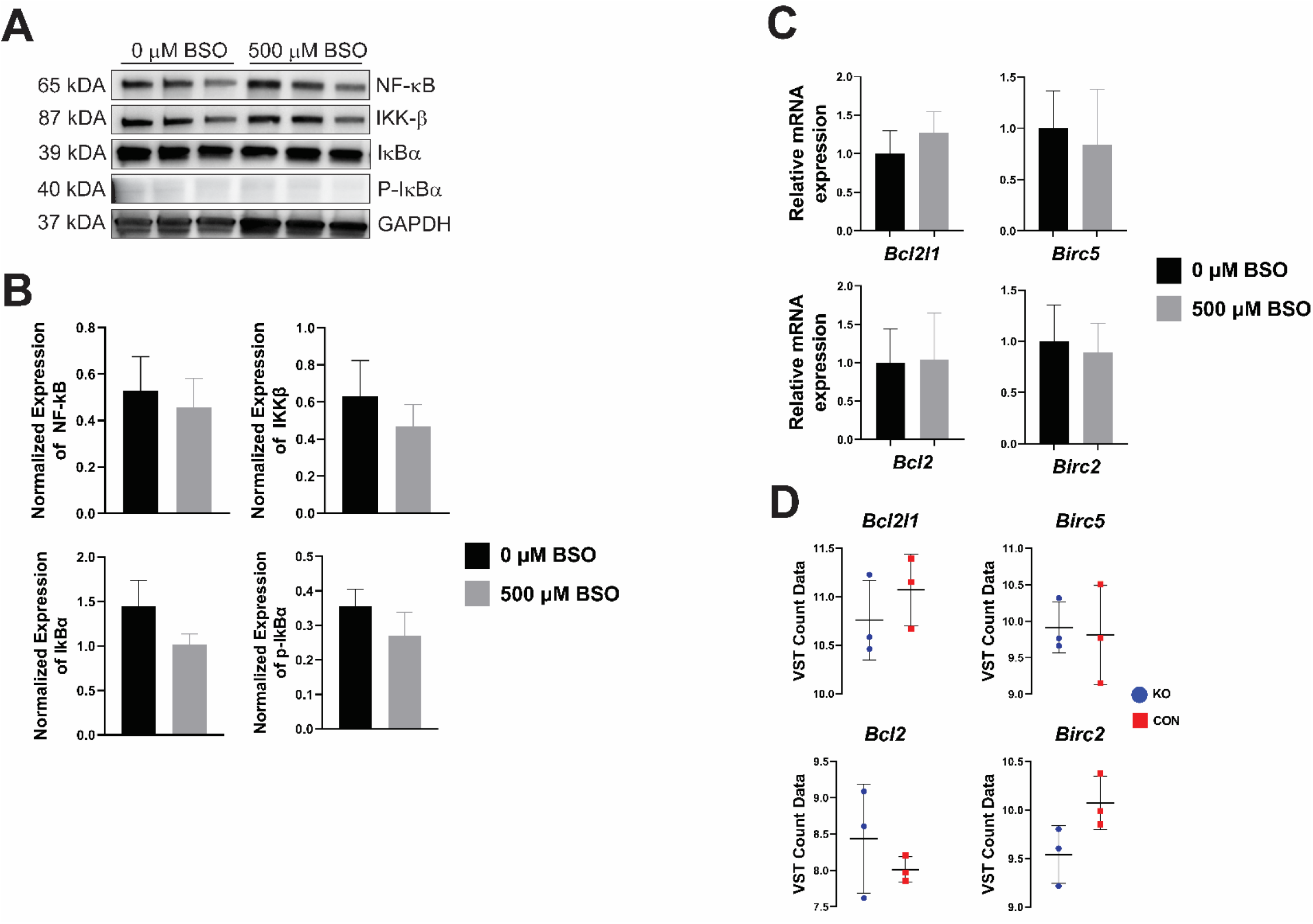
NF-κB activation in lens epithelial cells and lenses. 21EM15 cells were treated for 48 hours with 500 μM BSO and analyzed for NF-κB activation by quantifying the expression and activation (phosphorylation, P) of members of the NF-κB signaling pathway. (**A**) Representative Western blot of control (0 μM BSO) and treated (500 μM BSO) cells. (**B**) NF-κB signaling pathway protein (Ikk-β, NF-κβ, IκBα) and phosphorylated protein (P-IκBα) expression in control and BSO-treated cells were normalized to Ponceau S stain. Data are presented as the mean and associated standard deviation from three independent experiments. No differences occurred between BSO-treated and control cells (Student’s unpaired t-test, with P < 0.05 being considered significant). (**C**) Expression of genetic targets of NF-κB in control (0 μM BSO) and treated (500 μM BSO) 21EM15 cells. Gene expression was determined by reverse transcriptase (RT) quantitative PCR (qPCR) with the ΔCt method and GAPDH used as an internal normalization control. Gene expression is presented as the mean fold-change relative to control with associated standard deviation from three independent experiments. No differences occurred between BSO-treated and control cells (Student’s unpaired t-test, with P < 0.05 being considered significant). (**D**) RNA-seq box plots indicating variance stabilizing transformed (VST) normalized count data for genetic targets of NF-κB in the lenses of KO mice compared with the lenses of CON mice aged P1. VST count data are shown as mean (thin horizontal bar) ± standard deviation (error bar). Significance was evaluated at P < 0.05 and determined by the Benjamini-Hochberg method.

**Supplemental Figure 5:**
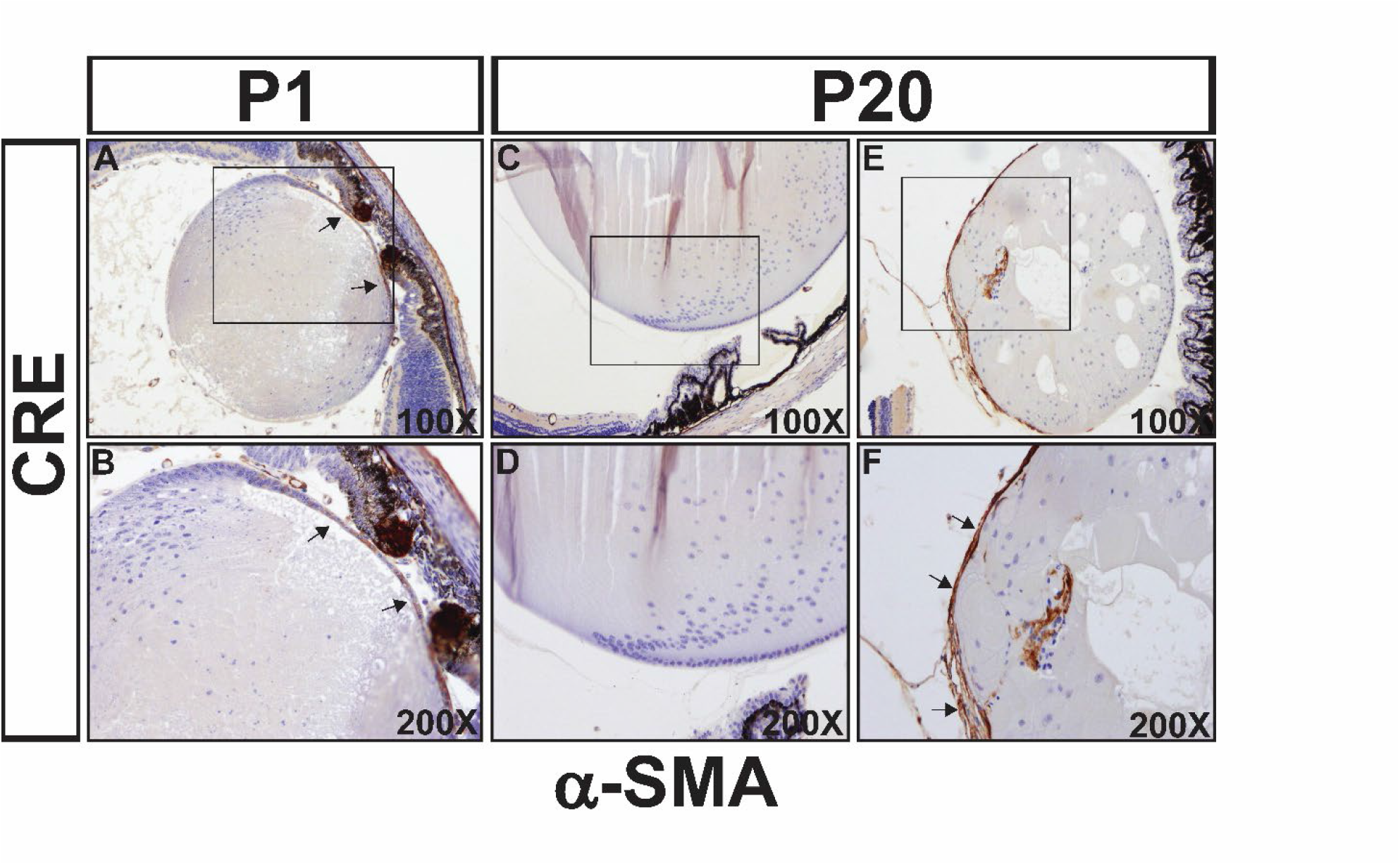
Analysis of a marker of EMT in the lenses of CRE mice aged P1 and P21. Immunohistochemical staining for α-SMA (closed arrows) and counterstaining with hematoxylin in the eyes of CRE mice aged postnatal day (P) 1 (A, B) and 21 (C-F). Regions demarcated by boxes in A, C and E are shown in higher magnification in B, D and F, respectively. Magnification is indicated in the lower right corner of each image.

**Supplemental Figure 6:**
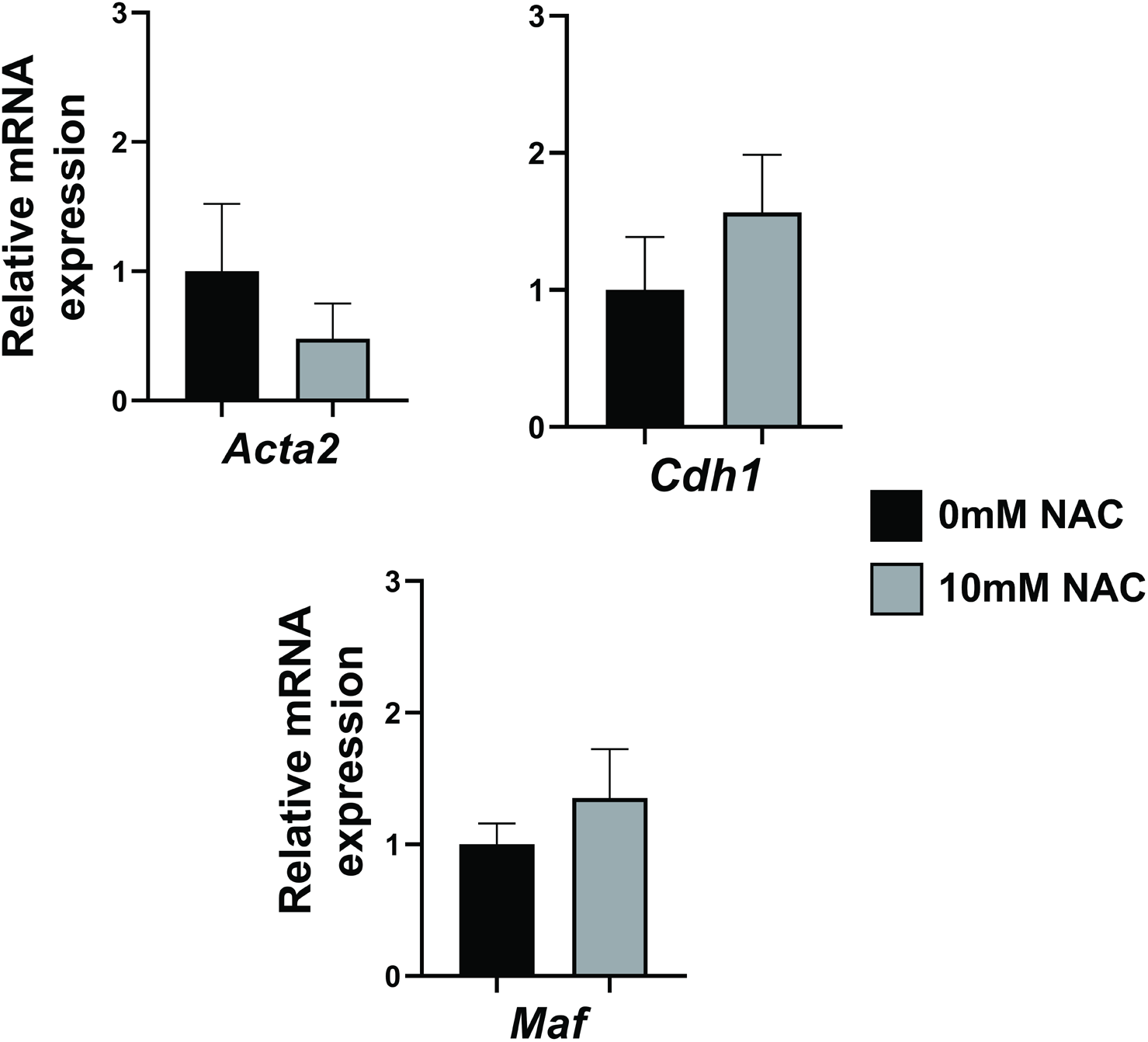
Analysis of EMT-related genes in lens epithelial explants. Lens epithelial explant systems were established in normal media or media containing 10 mM *N-*Acetyl-L-cysteine (NAC) and cultured for 24 hours. Gene expression was determined by reverse transcriptase (RT) quantitative PCR (qPCR), as calculated by the ΔCt method. GAPDH was used as an internal normalization control. Gene expression is displayed as the average of the fold change relative to control and associated standard deviation. * P-value < 0.05 Student’s unpaired t-test, compared to untreated (0 mM) explants.

**Supplemental Figure 7:**
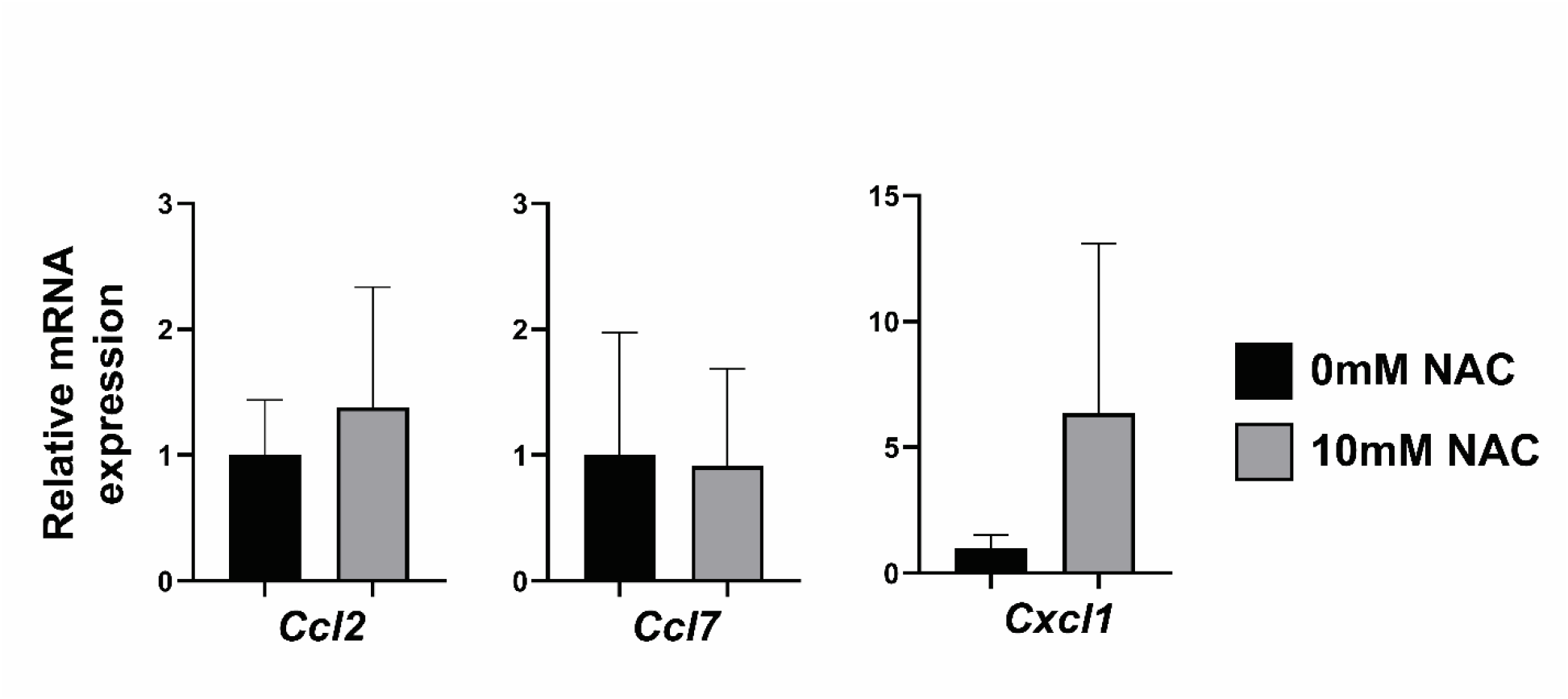
Analysis of cytokine expression in lens epithelial explants. Lens epithelial explant systems were established in normal media (0mM NAC) or media containing 10 mM *N-*Acetyl-L-cysteine (10mM NAC) and cultured for 24 hours. Gene expression was determined by reverse transcriptase (RT) quantitative PCR (qPCR), as calculated by the ΔCt method. GAPDH was used as an internal normalization control. Gene expression is displayed as the average of the fold-change relative to control, with associated standard deviation. * P < 0.05. Student’s unpaired t-test, compared to untreated (0 mM) explants.

**Supplemental Table 1:**
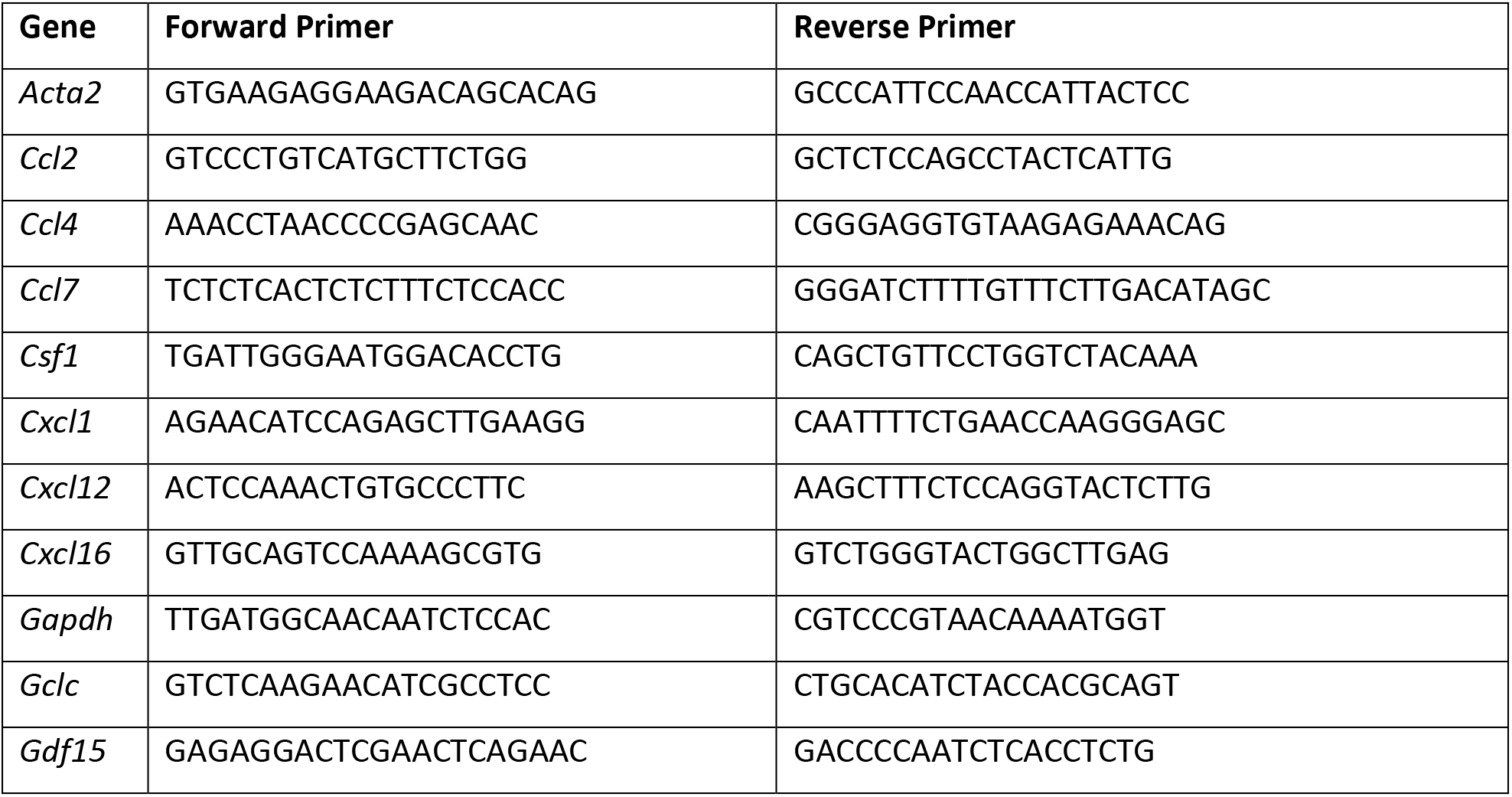
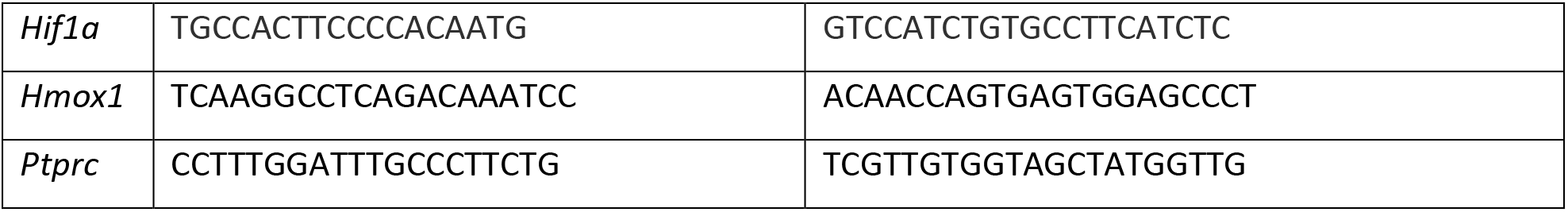
Primers used for RT-qPCR.

**Supplemental Table 2:** Top 25 upregulated genes in the lenses of KO mice aged P1 compared with lenses from similarly aged CON mice.

| Gene Symbol | log2FC<sup>a</sup> | adjusted P value<sup>b</sup> |
| --- | --- | --- |
| Arg1 | 7.411177 | 2.10E-56 |
| Ptgs2 | 7.235448 | 2.57E-38 |
| 3300005D01Rik | 6.342014 | 1.10E-25 |
| Cdsn | 6.030009 | 7.87E-22 |
| Slc14a1 | 6.014576 | 2.95E-22 |
| Ccl7 | 5.991658 | 8.79E-83 |
| Rbp4 | 5.704899 | 3.70E-28 |
| Cdkn1a | 5.680406 | 3.39E-57 |
| Dglucy | 5.623126 | 1.75E-18 |
| Arr3 | 5.587417 | 7.37E-22 |
| Ccl4 | 5.574396 | 3.39E-18 |
| Ccdc141 | 5.544726 | 1.31E-19 |
| Ch25h | 5.454229 | 3.18E-16 |
| Gdf15 | 5.42999 | 1.97E-26 |
| Cd300ld | 5.392139 | 1.97E-16 |
| Car2 | 5.338622 | 1.52E-70 |
| Cxcl16 | 5.274513 | 1.15E-15 |
| Clrn2 | 5.215495 | 3.66E-18 |
| Fgl2 | 5.134096 | 1.78E-15 |
| Tns4 | 5.082742 | 3.80E-14 |
| Adrb2 | 5.044112 | 2.34E-40 |
| Cd300lb | 5.007534 | 2.49E-13 |
| S1pr2 | 4.960696 | 2.49E-13 |
| Slc37a2 | 4.801622 | 4.07E-12 |
| Dusp10 | 4.792521 | 8.03E-13 |

<sup>a</sup> log2FC = KO compared with CON
<sup>b</sup> adjusted P value = Benjamini-Hochberg method

